# Red cabbage juice-mediated gut microbiota modulation improves intestinal epithelial homeostasis and ameliorates colitis

**DOI:** 10.1101/2023.08.23.554560

**Authors:** Emily Jean Wilson, Nagabhishek Sirpu Natesh, Parsa Ghadermazi, Ramesh Pothuraju, Marudhupandiyan Shanmugam, Dipakkumar R. Prajapati, Sanjit Pandey, Jussuf T. Kaifi, John R. Dodam, Jeffrey Bryan, Christian L. Lorson, Aude A. Watrelot, Jason M. Foster, Thomas J. Mansel, Siu Hung Joshua Chan, Surinder K. Batra, Jeyamkondan Subbiah, Satyanarayana Rachagani

## Abstract

Gut microbiota plays a crucial role in inflammatory bowel disease (IBD) and has therapeutic benefits. Thus, targeting the gut microbiota is a promising therapeutic approach for IBD treatment. We recently found that red cabbage juice (RCJ) ameliorates dextran sulfate sodium (DSS)-induced colitis in mice. However, the underlying mechanisms remain unknown. The current study investigated the modulation of gut microbiota in response to treatment with RCJ to ameliorate the DSS colitis. The initial results demonstrated that mice treated with DSS + RCJ showed increased body weight and decreased diarrhea and blood in feces compared to the DSS alone group. RCJ ameliorated colitis by regulating the intestinal barrier function by reducing the number of apoptotic cells, improving colonic protective mucin, and increasing tight junction protein in RCJ + DSS groups compared to the DSS group. Short-gun metagenomic analysis revealed significant enrichment of short-chain fatty acid (SCFAs)-producing bacteria *(Butyrivibrio, Ruminococcaceae, Acetatifactor muris, Rosburia Sp.* CAG:303*, Dorea Sp.* 5-2) increased PPAR-© activation, leading to repression of the nuclear factor κB (NFκB) signaling pathway, thus decreasing the production of crucial inflammatory cytokines and chemokines in the RCJ + DSS groups compared to the DSS group. Pathway abundance analysis showed an increased abundance of the SCFA pathway, reduced histidine degradation (*Bacteroides sartorii, and Bacteroides caecimuris*), and LCFA production in the RCJ+DSS treated group, suggesting the promotion of good colonic health. Furthermore, increased T-reg (FOXP3+) cells in the colon were due to SCFAs produced by the gut microbiota, which was corroborated by an increase in IL-10, a vital anti-inflammatory cytokine. Thus, our study provides the first evidence that RCJ ameliorates colonic inflammation by modulating the gut microbiota.

## Introduction

Inflammatory bowel diseases (IBDs), including Crohn’s disease and ulcerative colitis, are two major pathological entities that are a significant health problem in most developed and developing countries. In 2015, the Centers for Disease Control and Prevention (CDC) estimated that 1.3% of all adults in the USA, accounting for a total of 3.1 million patients, and 1 in 123 people in the UK suffered from IBD. Worldwide, at least 10 million IBD cases have been reported (1).

The primary clinical symptoms of IBD include acute abdominal pain, rectal bleeding, weight loss, anemia, hematochezia, diarrhea, negative effects on the immune system, and mortality risk (2). Although the complete pathogenesis of IBD is yet to be fully understood, impairment of the epithelial cellular barrier, inflammation-mediated immune dysfunction, and dysbiosis of the gut microbiota are hallmarks of IBD (3). It has been proposed to be associated with the host’s genetic susceptibility, alterations in intestinal microbiota, environmental factors, and immunological abnormalities (4). Typically, intestinal epithelial barrier disruption leads to the translocation of microbiota from the lumen to the lamina propria, eventually triggers inflammation (5).

Current IBD treatments include anti-inflammatory drugs, such as corticosteroids, immunosuppressors (TNF-α inhibitors), antibiotics, and surgery (6). However, there are significant risks associated with these treatments; for example, tumor necrosis factor-alpha (TNF-α) inhibitors can increase the risk of infections, leading to illnesses such as tuberculosis (7). Additionally, the long-term use of antibiotics, including ciprofloxacin and metronidazole, is widespread in patients with IBD, which can lead to antibiotic resistance and negatively alter the gut microbiota (8). Due to therapy-related toxicity risks and the ineffectiveness of currently administered drugs for IBD, there is an urgent need to seek an alternative and effective therapeutic approach to treat IBD at initial diagnosis and for the high burden of recurrence.

Recent studies have highlighted that alterations in the gut microbiota (dysbiosis) play a vital role in IBD pathogenesis (9–11). In this context, research efforts have been directed to understand the role of gut microbiota alteration in pathogenesis, aiming to restore the gut microbiome composition using various nutraceuticals (prebiotics), probiotics, symbiotics, and fecal microbiota transplantation as an alternative therapy to ameliorate intestinal inflammation in IBD. Efforts to develop alternative strategies, such as using nutraceuticals that can improve the efficacy and safety of existing therapies, are urgently needed to improve the quality of life of IBD patients.

Nutraceuticals can treat IBD directly as they contain bioactive compounds, such as secondary metabolites, polyphenols, and poly/oligosaccharides (carbohydrates indigestible by host digestive enzymes) from plants that are beneficial to the intestinal epithelium. These polyphenols (anthocyanins) have been shown to have direct anti-inflammatory and antioxidant effects on the intestinal epithelium, thus acting as therapeutics for IBD patients (12). Moreover, nutraceuticals can indirectly treat IBD by modulating the gut microbiota (13). The microbiome converts these compounds into other compounds with altered biochemical functions that are absorbed through the intestinal epithelium. However, only a certain taxonomic group has been associated with this microbial conversion, which is barely understood at the genetic and biochemical levels (14).

In the present study, red cabbage (*Brassica oleracea L. var. capitata f. rubra*) juice extract, which is widely used in traditional medicine because of its rich minerals, oligosaccharides, and several bioactive substances such as glucosinolates (GSLs), indole–sulfur phytoalexins, S-methylmethionine, and phenolic compounds such as flavonol glycosides, acylated anthocyanins, and hydroxycinnamic acid derivatives (15, 16). Anthocyanins, major polyphenol pigments, have been reported to reduce acute and chronic colitis in mouse models (17). GSLs, hydrolyzed by myrosinase, produce bioactive compounds, such as indoles and isothiocyanates (ITCs) (18). In recent studies, bioactive compounds of *Brassica* species have been associated with the modulation of the gut microbiome (19).

The aim of the current study was to determine the nutraceutical potential of RCJ to investigate whether it can ameliorate IBD through modulation of the gut microbiota in DSS induced colitis model.

## Methods

### Pulsed Electric Field-assisted extraction (PEF) of red cabbage juice (RCJ)

Red cabbages were obtained from the local market and processed by using PEF with the processing parameters (1.2 kV/cm, 2 μF, 25 pulses, 3.43 kJ/kg), The samples of mash derived from PEF-treated red cabbages were mechanically pressed (450 N, 9 min). The extract of red cabbage juice (RCJ) stored at -80 °C and used for further biochemical assays. The pH was measured using a pH meter.

#### Phenolic compounds quantification of RCJ

The initial bioactivity of PEF-treated RCJ was analyzed for total phenolic compounds and total anthocyanin concentrations. An aliquot of each of the sample frozen at -80°C was used for to determination of the same bioactivity components to test the effects of freezing. The total iron-reactive phenolic compound concentration in RCJ was quantified by UV-Vis spectrophotometry using a previously published method by UV-Vis spectrophotometry (20). RCJ anthocyanin concentration was analyzed using a 375 1260 Infinity II HPLC (Agilent Technologies, Santa Clara, CA) with a reserved-phase column (Li Chrospher 100-5 RP18 250 × 4.0 mm, Agilent Technologies), DAD (Agilent 1260 377 Infinity II DAD WR) and fluorescence detector (FLD) (Agilent 1260 Infinity II FLD Spectra) as previous reports (20). Ammonium dihydrogen phosphate (50 mM) was used as the mobile phase at pH 2.6 (mobile phase A), 20% (v/v) mobile phase A in acetonitrile (mobile phase B), 0.2 M ortho-phosphoric acid in water, pH 1.5 (mobile phase C). The temperature of the column was maintained at 40 °C at a flow rate of 0.5 mL/min. The sample supernatant (20 μL) was injected. Monomeric phenolics were identified and quantified at different wavelengths:280 nm for gallic acid, 360 nm for flavonols, 316 nm for hydroxycinnamic acids, and 520 nm 385 for anthocyanins. FLD was used to detect and quantify (+)-catechin and (−)-epicatechin at an excitation wavelength of 276 nm and an emission wavelength of 316 nm. cyanidin-3-O-glucoside was used as a standard to quantify anthocyanins present in RCJ.

#### Sugars and Organic acids were quantified in RCJ

Sugars and Organic acids were quantified in the RCJ using high-performance liquid chromatography (HPLC) (1200 series, Agilent Technologies) with a diode-array detector (DAD) and refractive index detector (RID) (Agilent 1200 series). Bio-Rad Aminex HPX-87H and Bio-Rad fermentation monitoring columns with H+ guard cartridges were used, and a sample of 10 μL was injected. Sulfuric acid (5 mM) was used as the mobile phase with a flow rate of 0.65 mL/min for 35 min. Malic acid detection was performed at 210 nm using DAD. Other residual sugars were detected by using RID (cell temperature of 55 °C). Commercial standards from Bio-Rad were used to obtain calibration curves for each compound.

### DSS-induced colitis

To evaluate the protective role of RCJ in DSS-induced colitis, C57/BL/6J mice (male and female) were divided into two groups. The first group (N=20 mice) served as a control, and phosphate-buffered saline (PBS) was administered. In the second group (N=30 mice), RCJ (200 µL) was administered per day by oral gavage throughout the experiment. All mice (50 total) were fed *adlibtum*. After eight weeks, both groups were further divided into two subgroups: PBS (N=5) and DSS (N=15) from the first group, and RCJ (N=15) and DSS + RCJ (N=15) from the second group. As described previously, colitis was induced by administering two cycles of 3% (w/v) DSS in drinking water (21). There was one week of DSS, followed by one week of recovery without DSS and a second week of DSS. RCJ and PBS oral gavage administration was continued for these three weeks. Mice were euthanized by CO_2_ asphyxiation, followed by cervical dislocation at the end of the experiment. The blood and other organs were collected and stored.

### Disease severity

Diarrhea, blood in feces scores, and mouse body weight were monitored throughout the experiment. Diarrhea scores and blood in feces scores were recorded daily and scored from 0 to 3 (absent, mild, moderate, and severe). To assess disease severity, the disease activity index (DAI) was calculated, as previously described (22). Briefly, the DAI was calculated as a combination of weight loss, diarrhea, and bleeding scores, resulting in a colitis score of 0–10 (unaffected by severe colitis). Body weight scores were calculated based on the change in body weight from the original weight (≥0%=0; -5% to 0%=1; -10% to -6%=2; -15% to -11%=3; and <-15%=4). This bodyweight score was combined with diarrhea and blood in feces scores to calculate DAI.

### Cytokine analysis

Pro-inflammatory cytokines, such as IFN-γ, TNF-α, IL-1α, IL-1β, IL-1ra, IL-5, IL-6, IL-10, IL-12, IL-13, IL-17, and IL-23, are associated with IBD (23, 24). These cytokines were analyzed using a cytokine array kit according to the manufacturer’s protocol (Proteome Profiler Mouse Cytokine Array Kit, Panel A; R&D Systems, Minneapolis, MN, USA). Plasma samples (100 μL/group) were pooled within each group, with five samples in the PBS group and six in the other groups. The results were quantified by analyzing the pixel density of each spot on the developed film using the ImageJ software (http://rsb.info.nih.gov/ij).

### Histology

Colon tissue samples harvested from the mice were fixed in 10% buffered formalin. The samples were then processed and embedded in paraffin blocks. From each block, 4 μm sections were prepared. The prepared colon tissue sections were stained with hematoxylin and eosin (H&E) and periodic acid-Schiff (PAS) for all treatment groups (N=6/group). The H&E-stained tissues were scored based on inflammatory cell infiltration (0-3) and changes in intestinal architecture change (0-3). These two scores were summed to obtain a final score of 6. The PAS-stained tissues were analyzed by counting the number of PAS-positive cells. The number of cells per crypt is reported, with a minimum of 20 crypts in each section.

### Terminal deoxynucleotidyl transferase dUTP nick end labeling (TUNEL) assay

A commercially available kit (Abcam, ab206386) was used to detect apoptotic cells in mouse pancreatic cancer tissues. Briefly, 5 mm thick tissue sections were deparaffinized in xylene and further dehydrated with a series of graded alcohol, followed by proteinase K treatment and 3% H_2_O_2_ treatment to inactivate endogenous peroxidases in the cells. Biotin-labeled deoxynucleotide incorporation in apoptotic cells, catalyzed by terminal deoxynucleotidyl transferase (TdT), was detected by incubation with streptavidin-horseradish peroxidase (HRP) conjugate. Signals were detected using 3,3′-diaminobenzidine (DAB) substrate, and sections were counterstained with methyl green. Positive and negative control tissues, treated with DNase I and water instead of TdT, were used for comparison.

### Immunohistochemistry/immunofluorescence

For immunohistological analysis, the prepared tissue sections were heated overnight and then deparaffinized in xylene (2x, 5 min). The sections were then hydrated using graded alcohol. Then, citrate buffer (0.01M, 95°C, pH 6.0) was used for antigen retrieval. 0.3% H_2_O_2_ in methanol for 30 min was used to quench the endogenous peroxidase activity. After PBS washing, sections were blocked with 2.5% horse serum (ImmPRESS kit; Vector Labs, Burlingame, CA, USA) for 2 h. Next, sections were incubated with primary antibodies and kept at 4°C overnight. After washing with PBS containing 0.01% Tween 20 (PBST), the sections were incubated for 30 min with a secondary antibody (peroxidase-labeled anti-mouse/anti-rabbit IgG (ImmPRESS kit, Vector Labs, Burlingame, CA, USA)). The sections were washed with PBST (3x, 5 min) and developed using a DAB substrate kit (Vector Laboratories, Burlingame, CA, USA). Counterstaining was performed with hematoxylin. Graded alcohol was used to dehydrate the slides, followed by xylene (2x, 5 min). After drying, the slides were mounted in toluene and imaged (25, 26). The following antibodies from various vendors were used for the immunohistochemistry study: anti-Ki 67 (Abcam, ab15580); Anti-SOD1 (CST, 65778); Anti-GPX4 (Abcam, ab125066); Anti-4-Hydroxynonenal (R&D Systems; Biotechne, MAB3249); Anti-MUC2 (Abcam, ab272692); Anti-MUC4 (Santa Cruz, SC-33654); anti-claudin 1 (Proteintech, 13050-1-AP); anti-occludin (Proteintech, 66378-1-Ig); Anti-F4/80 (Abcam, ab300421); anti-pIKKβ (CST, 36214SF); anti-pNF-kB (Abcam, ab131100); anti-TNF-α (Abcam, ab1793); Anti-pSTAT3 (CST, 9145); Anti-CD3 (CST, 78588); Anti-RORγ (Abcam, ab207082); Anti-FOXp3 (CST, 12653); Anti-MPO (Proteintech, 22225-1-AP); Anti-Ppar γ (Proteintech, 16643-1-AP). All immunostained slides were analyzed by a pathologist. Slides were assessed based on a previously reported (27). For immunofluorescence staining, antibodies against ZO-1 (Proteintech, 21773-1-AP), iNOS (Proteintech, 22226-1-AP), and COX-2 (Proteintech, 66351-1-Ig) were used. Appropriate secondary antibodies conjugated to Alexa Fluor 488 or 594 (Vector Laboratories) were added at 1:200 dilution, and nuclei were counterstained with 4’,6-diamidino-2-phenylindole (DAPI) (Vector Laboratories H-1800).

### Metagenomic sequencing

Mouse fecal samples were collected before and at the end of the study from all groups and stored at -80°C until fecal microbial DNA isolation. Microbial DNA was isolated from mouse fecal samples (QIAamp PowerFecal Pro DNA Kit; Qiagen, Hilden, Germany). At the end of the experiment, samples were analyzed from three animals in each group. The DNA samples were purified using Zymo Spin columns (Zymo Research). Libraries were prepared using a Nextera Flex DNA Kit (Illumina). The concentrations of the libraries were measured in a Qubit30 using a high-sensitivity kit. The quality (size distribution) of the libraries was assessed using an Agilent 2100 Bioanalyzer. The libraries were pooled at an equimolar ratio and denatured in the presence of NaOH. The loading concentration was 1.5 pm, and sequencing was performed in Nextseq NS500. A 150-base paired-end run and high-output flow cell were used. Sequencing was performed using Basespace (Illumina).

### Taxonomic and functional analysis

FASTQ files for each sample were aligned against the full NCBI NR database reference genome using the Diamond (28). The alignment files were processed using MEGAN 6 (29). Taxonomic abundances, which were generated based on the output from the MEGAN analysis, are represented in bar graphs. The groups were compared using the linear discriminant analysis (LDA) effect size (LEfSe) method (30). An LDA score of >2 was used to identify features that significantly discriminated among groups. A cladogram was generated based on differential abundance values using LEfSe. Box plots were generated using Statistical Analysis of Metagenomic Profiles (STAMP) analysis (31). The YAMP pipeline V 0.9.5.3 (32) was used to profile the metabolic functions of the samples, directly mapping the short reads to the reference databases of HUMAnN 3 and MetaPhlAn 3. YAMP leverages FastQC (33) and MultiQC (34) for quality check, BBDuk (35) for trimming, BBWrap for decontamination, MetaPhlAn (36) for taxonomic profiling, and HUMAnN (36) for functional profiling of metagenomic raw reads. Raw reads were decontaminated by removing reads mapped to the mouse genome with more than 97% similarity. Reads shorter than 70 bp were excluded. Additionally, the YAMP database of sequencing artifacts and adapters was used to trim the short reads using BBwrap. Trimmed reads were quality checked, and all samples showed satisfactory quality and number of reads in each sample. HUMAnN 3 in the YAMP pipeline was used for the functional profiling of quality-checked reads. The outputs of HUMAnN were normalized to count per million (CPM). Multivariable association discovery in population-scale meta-omics studies (MaAsLin2) (37) was used to extract the association between pathways and gene counts with different treatments (PBS, RCE, DSS, and RCE+DSS) using a linear mixed model. All statistics of pathway association with the treatment groups were extracted from the outputs of MaAsLin2. Any pathways or reactions associated with the sex of the mice were discarded from the final analysis. The significant results were filtered to exclude results with *q-value*>0.05, or *p-value*>0.05. Non-metric multidimensional scaling (NMDS) (38) in the Vegan R package v2.6-4 (39) was used to map pathway and taxonomy data to 2-dimensional space using. Non-metric multidimensional scaling (NMDS) plots were generated using the ggplot2 R package (40). The vegan package in R was used to create alpha diversity measures for each sample. MultiQC output, the raw outputs and log files of YAMP, YAMP configuration files that were used, and a Jupyter notebook providing the steps taken along with the parameters used in each step and the generated plots are all available at (https://github.com/chan-csu/RCJ_Megtagenomics).

#### Statistical analysis

Statistically significant differences were analyzed using the Student’s t-test with a 0.05 significance level (p<0.05). Linear mixed-effects models were used to analyze the changes in body weight and DAI scores over time. Treatment group, time, and treatment-by-time interactions were included in the model. The percent change in the body weight model was adjusted for the baseline weight. Pairwise comparisons were adjusted for multiple comparisons using Tukey’s method. The Kaplan-Meier method was used to estimate the survival curve from induced colitis, and survival times were calculated as the days from treatment initiation to death from colitis on the last study date. Animals that died from causes other than colitis were censored. Toxicity data were summarized descriptively over time using box plots. SAS software (version 9.4; SAS Institute Inc., Cary, NC, USA) was used for data analysis.

### Results

#### Bioactive compounds remain active despite freezing and PEF treatment

To understand whether PEF treatment, freezing, and thawing would significantly affect its biological activity, different biological parameters were tested, and it was observed that there was no substantial change in the bioactivity of RCJ (**Fig S table 1**). Furthermore, there was no significant change in total phenolic concentration or total monomeric anthocyanin content in RCJ (**Fig S table 2**).

Chemical parameter analysis was then carried out. Four red cabbages were procured (total weight, 3.8 kg), and 1.4 litres of juice was obtained using a food processor and filtered with a cheesecloth. The pH of RCJ was 6.42±0.05 with dissolved solids of approximately 6.4±0.2 gms. RCJ juice contained 18.5±0.2 gms of glucose, 15.1±0.2 gms of fructose, 0.8±0.1 gms of citric acid, 3±0.3 gms) malic acid with 34.2±2.1 gms of unknown acids per liter of RCJ **(SF 1A)**.

Spectrophotometry analysis of RCJ revealed that 382.5±93.5 mg of total phenolic compound and 257.7±3.1 mg of anthocyanins were present in 1 liter of RCJ (**Fig S table 2**). Next, HPLC-DAD was performed to determine the composition of polyphenols in RCJ; approximately eight monomeric polyphenols were identified in RCJ (**Fig S table 2**). In addition to polyphenols, RCJ also contains 254±23.6 and 55±2.3 mg of free anthocyanins, and total hydroxyl innamic acid per liter of red cabbage juice. After the characterization of organic acids and sugars in red cabbage juices, a high concentration of inorganic acid compounds was observed.

Because of the high fraction of the organic acid fraction, RCJ was further analyzed for glycosyl composition (sugar residue analysis) to quantify the monosaccharide composition of polysaccharides, oligosaccharides, or glycoproteins present in RCJ by GC-MS using TMS-derivatized monosaccharides. Our results revealed that the carbohydrates in RCJ are mainly composed of glucose residues, with galactose, fructose, arabinose, and mannose as the minor sugar components. (**Fig S1**). Detailed calculations of the molecular percentage of monosaccharides and the total carbohydrate percentage by weight for the RCJ sample are shown in (**S table 3**)

#### Prophylactic RCJ intervention alleviates DSS-induced colitis in mice

To directly assess the effect of RCJ on DSS-induced colitis development, C57BL/6 mice were divided into four groups: PBS, DSS, RCJ, and DSS+RCJ (**Fig 1A**. Interestingly, the DSS group showed a significant decrease in body weight (**Fig 1B**), colon length (**Fig 1C)** and higher blood and diarrhea scores (**Fig 1D & E**), whereas RCJ administration reverted back to normal. Furthermore, compared with the DSS group, RCJ supplementation resulted in a lower disease activity index (DAI) (**Fig 1F)**.

**Fig 1:**
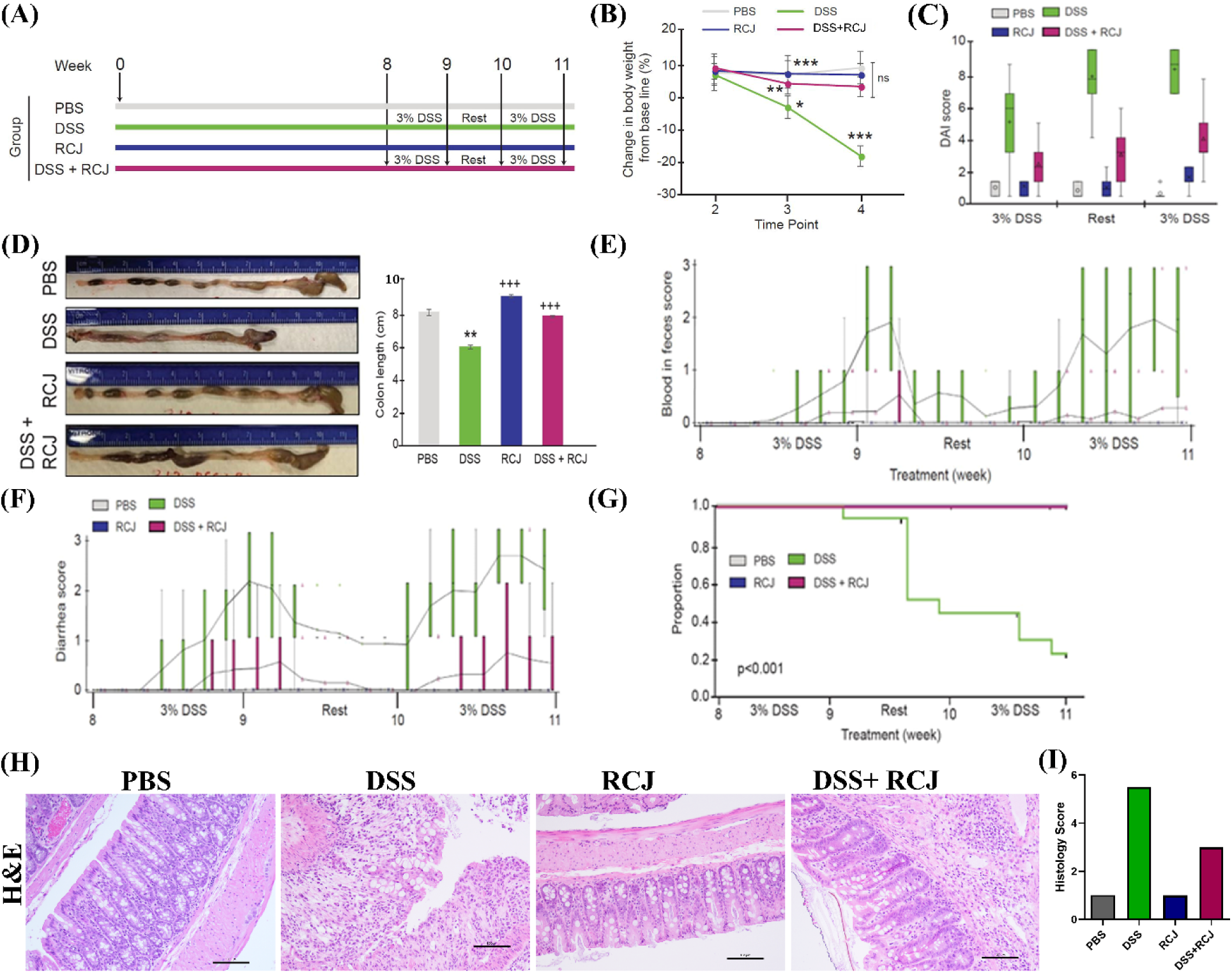
**(A)** Schematic diagram of the *in vivo* experimental design and access to water and standard feed. **(B)** Effect of DSS and RCJ on body weight where time point 1 is the baseline before treatments began. Time point 2 is during the RCJ treatments. Time points 3 is after the DSS administration began. Time points 4 is after a rest period. Error bars are the confidence interval (95%). Data were presented as mean±SEM (n=15 per group) **(C)** Effect of RCJ and DSS on colon length. Effect of RCJ and DSS on **(D)** kinetics of daily disease activity index (DAI) scores throughout the study duration. and **(E)** blood in feces scores. **(F)** Diarrhea scores **(G)** Survival curve (H) H&E-stained colon section. **(I)** histological scores of the colon (n=15 per group). Scale bars represent 100 µm. Statistical significance was determined using one-way ANOVA, followed by the Tukey test. *P ≤ 0.05, **P ≤ 0.01, ***P ≤ 0.001

Subsequently, the survival benefits were analyzed using RCJ in combination with DSS. The Kaplan-Meier Survival curve revealed a higher percentage of deaths due to severe colitis in mice treated with DSS than in the other groups (p<0.001) (**Fig 1G)**. No deaths were observed in the PBS, RCJ, or RCJ+DSS groups. Times were censored at the end of the study owing to deaths unrelated to colitis, such as misadministration of oral gavage.

In addition, H&E staining showed increased crypt depth, submucosal edema, macroscopic spaces between crypts, less hyperplasia of crypts, low epithelial cell damage, damaged brush borders (villi), reduced mononuclear cell infiltration in the submucosa, and reduced colon inflammation upon RCJ supplementation (**Fig 1H).** Moreover, an increase in the overall histological score (parameters for the histology score were 1. Architectural damage 2. The extent of inflammation was 3. Chronic inflammatory infiltrate) was observed in the RCJ group compared to that in the DSS alone group (**Fig 1I)**.

#### RCJ ameliorates colitis by regulating intestinal barrier function and DSS-induced oxidative stress in the intestinal mucosa

Healthy epithelial cells are critical for maintaining intestinal barrier function. Proliferation and apoptosis are two key factors in the differentiation of healthy epithelial cells. Colonic epithelial cell proliferation was assessed by Ki-67 staining, where the DSS+RCJ group showed a significantly higher number (p < 0.05) of Ki-67 positive cells when compared to the DSS group (**Fig 2A**). Furthermore, RCJ treatment reduced the number of TUNEL-positive nuclei in the colonic epithelium compared to that in the DSS group (**Fig 2A).** The DSS+RCJ group showed significantly reduced TUNEL-positive nuclei, indicating reduced apoptosis. Next, the influence of RCJ on oxidative stress, superoxide dismutase (SOD), 4-Hydroxynonenal (4-OH-enol), and glutathione peroxidase 4 (GPX4) in the colon tissue was measured. Compared with the DSS group, the DSS+RCJ group showed suppressed concentrations of SOD, GPX4, and 4-OH-enol. Compared with the PBS controls, colitis was attenuated mainly by RCJ (**Fig 2B)**. These results showed that RCJ treatment markedly ameliorated DSS-induced colitis. These results show that RCJ improved the integrity of the reduced intestinal permeability and intestinal epithelial barrier, thus restoring the intestinal barrier function.

**Fig 2:**
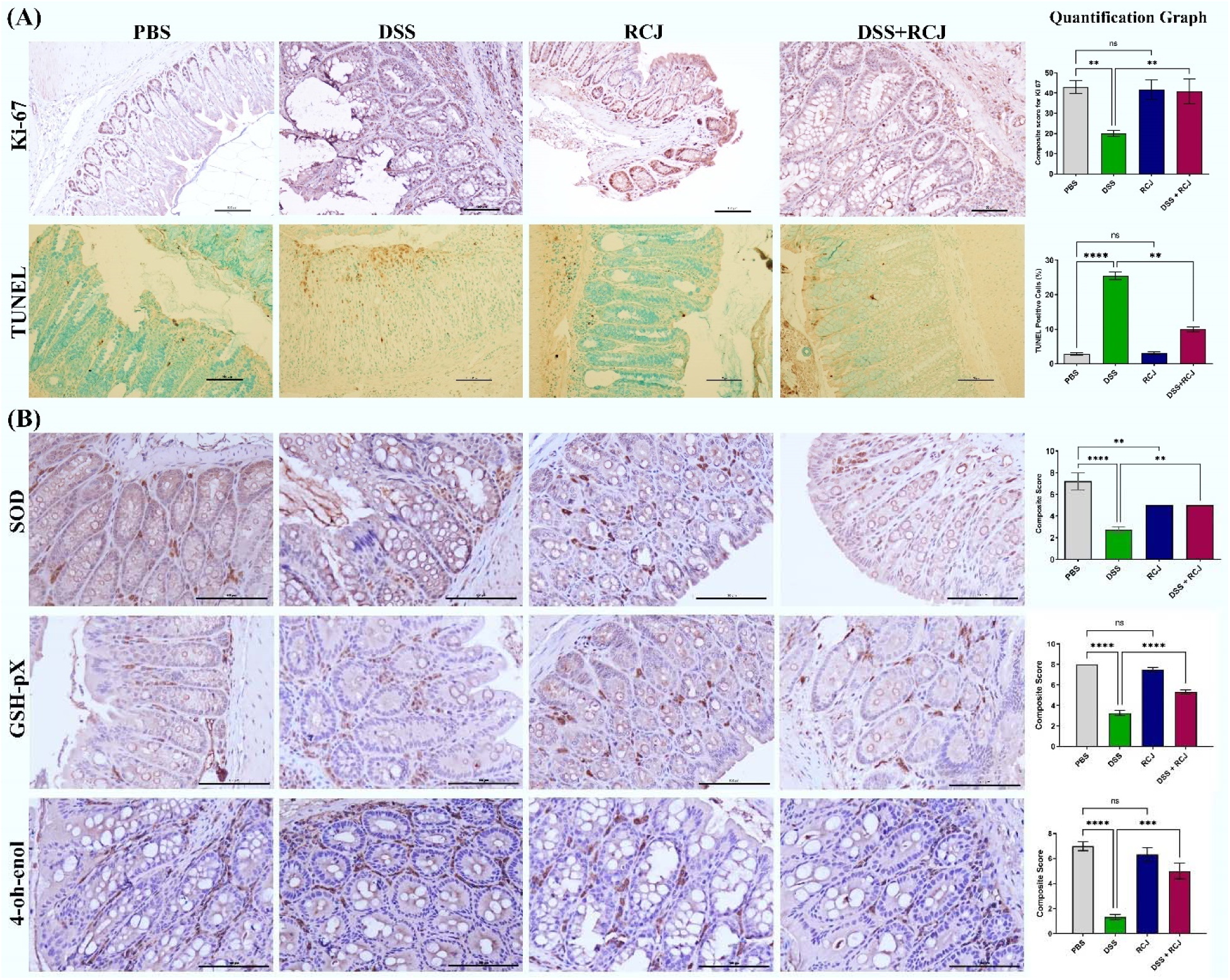
RCJ attenuated oxidative stress and colonic damage. **(A)** Depicts IHC staining for Ki67 marker in colonic epithelium staining to assess epithelial cell proliferation and TUNEL-positive nuclei (apoptotic cells) in the colonic epithelium in brown (**B)** Representative colons from each treatment group; scale is in cm. Error bars in the histograms are the standard error of the mean. Scale bars represent 100 µm. Statistical significance was determined using one-way ANOVA, followed by the Tukey test. *P ≤ 0.05, **P ≤ 0.01, ***P ≤ 0.001

Hematoxylin and eosin staining showed that the DSS group had intense, severe epithelial cell damage, submucosal edema, including shortening and hyperplasia of crypts, inflammatory lesions, macroscopic spaces between crypts, and severe inflammatory cellular infiltration in the submucosa. RCJ treatment notably ameliorated the colon inflammatory symptoms, including less inflammatory cell infiltration, relative submucosal edema, intact surface epithelium, normal crypt glands, and mild submucosal edema, and the condition of the colon was close to that of PBS mice. The DSS-treated group showed a significant reduction in the thickness of the colonic epithelial mucosa, which was attenuated to near normal levels. Thus, to assess the effect of RCJ on the colonic mucosal barrier, Alcian blue and PAS staining were used to check mucin-secreting goblet cells in the colonic epithelia (**Fig 3A**). Next, intestinal homeostatic mucins (MUC2 and MUC4) and supplementation with RCJ resulted in a significant (p<0.001) increase in the expression of colonic MUC2 and MUC4 (**Fig 3B).** Furthermore, RCJ-treated animals showed improved intestinal barrier function and tight junction proteins, such as ZO-1, Occludin, and Claudin-1 (**Fig 3C & D**).

**Fig 3:**
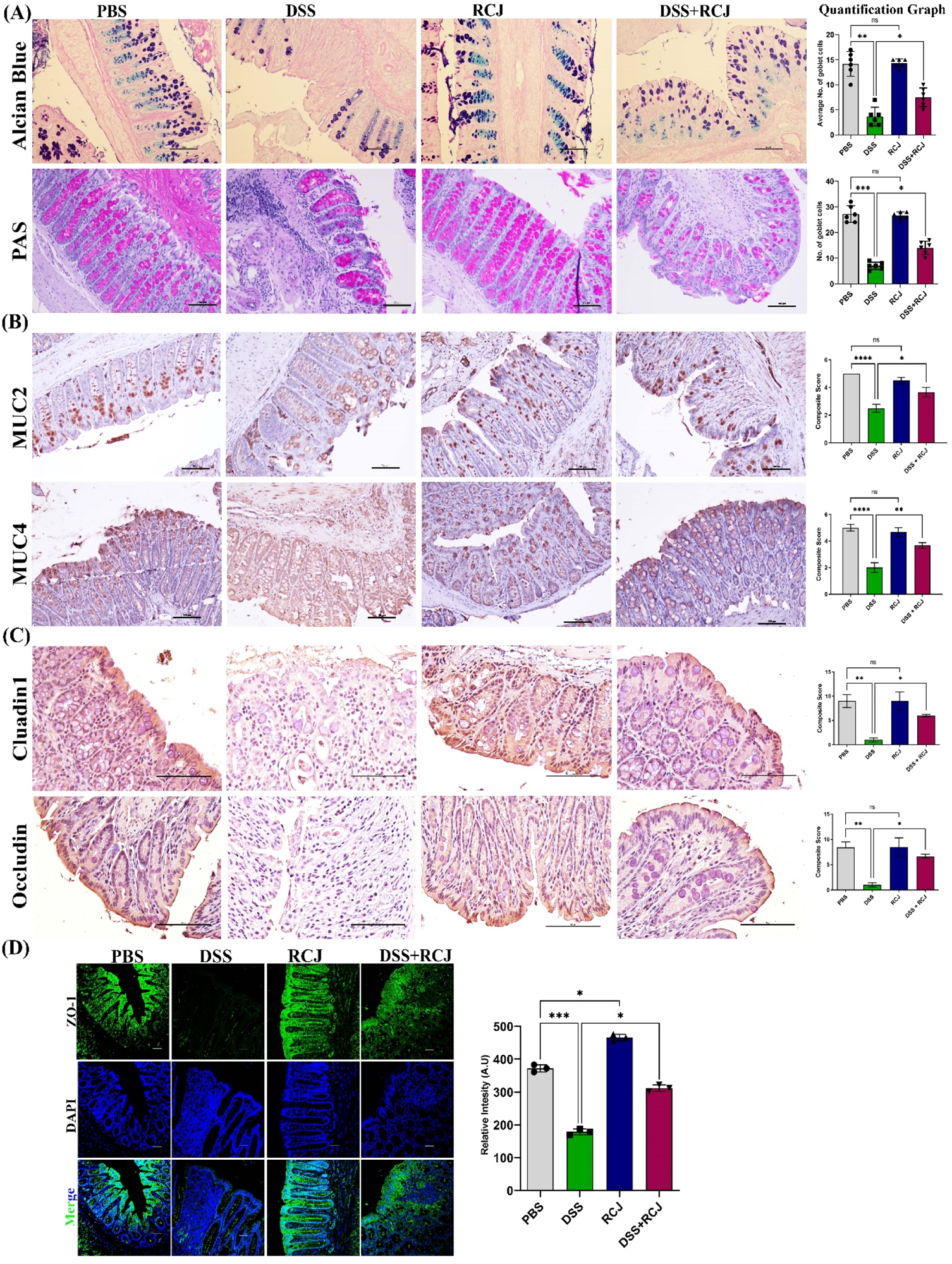
RCJ ameliorates colitis by regulating the intestinal barrier function. **(A)** Represents Alcian blue staining denoting the mucin-secreting goblet cells in the colonic epithelia and PAS-positive cells denoting acid and neutral mucin **(B)** Depicts IHC staining for MUC2 and MUC4, which was stained to understand the protective mucin layer **(C)** Shows IHC staining for tight junction markers claudin and Occludin **(D)** Immunostaining for colonic ZO-1, (green) counterstained with the nuclear stain DAPI (blue). (**E**) Shows relative intensity, quantitative analysis of the fluorescence by Image J. All values represent the means ± SD; error bars in the histograms are the standard error of the mean. Scale bars represent 100 µm for the IHC and 50 µm for IF. Statistical significance was determined using one-way ANOVA, followed by the Tukey test. *P ≤ 0.05, **P ≤ 0.01, ***P ≤ 0.001

#### Prophylactic RCJ intervention alleviated colonic pro-inflammatory status

To elucidate how RCJ reduced colitis severity, the expression of pro- and anti-inflammatory cytokines and chemokines was quantified using a cytokine array (IL-1α, IL-1β, IL-3, IL-6, IL-10, IL-16, IL-17, IL-23, IL-27, CCL1, CXCL-1, CXCL-9, CXCL-10, CXCL-11, G-CSF, GM-CSF, TNF-α, and IFN-γ). DSS-treated mice had elevated levels of crucial pro-inflammatory cytokines such as TNF-α, IFN-γ, IL-1β, and IL-6, and chemokines such as CXCL1 and CXCL 9-11. In contrast, RCJ treatment resulted in significantly (p>0.005) lower levels of various pro-inflammatory cytokines and chemokines and elevated levels of the anti-inflammatory cytokine IL-10 (**Fig 4A-4L**). Furthermore, iNOS and COX 2 were also reduced by RCJ treatment (p < 0.05 & p<0.001, respectively) (**Fig 4M)**. These results suggest that RCJ can diminish the pro-inflammatory cytokine response in DSS-induced colitis in mice.

**Fig 4:**
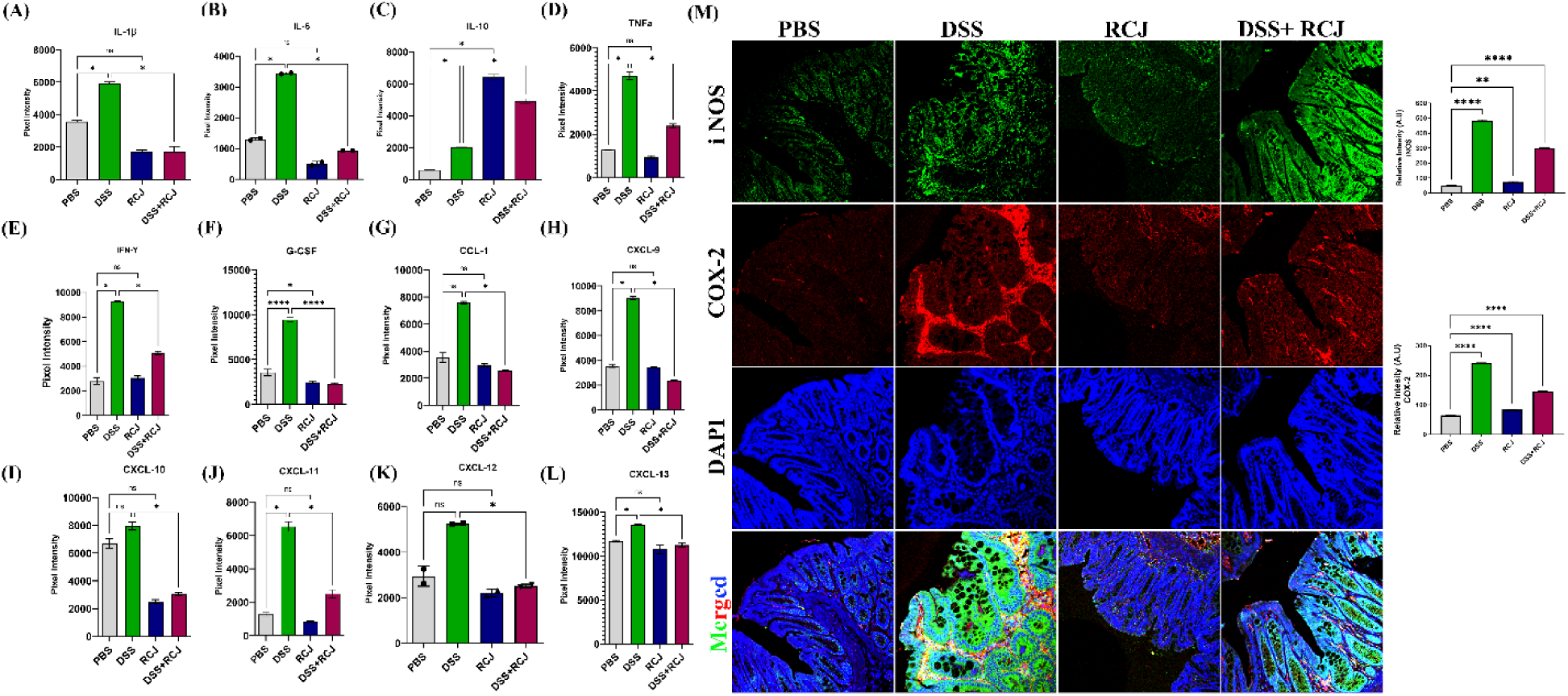
To understand the effect of RCJ on colonic pro-inflammatory status **(A-L),** Graphs show **a** quantified expression of pro- and anti-inflammatory cytokines and chemokines using a cytokine array **(M)** Immunofluorescence staining for iNOS (green), COX-2 (red) and counterstained with the nuclear stain DAPI (blue). Graphs show relative intensity, and quantitative analysis of the fluorescence by Image J. All values represent the means ± SD; error bars in the histograms are the standard error of the mean. Scale bars represent 50 µm for IF. Statistical significance was determined using one-way ANOVA, followed by the Tukey test. *P ≤ 0.05, **P ≤ 0.01, ***P ≤ 0.001

#### Prophylactic RCJ intervention restores microbiome diversity and content

The release of inflammatory cytokines and chemokines, which could be due to an abnormal gut microbiota composition, would induce an altered immune response, including the release of inflammatory factors and aggregation of inflammatory cells. Our biochemical analysis showed that the presence of oligo- and polysaccharides in RCJ could modulate gut microbiota composition. To address this, shotgun metagenomic sequencing was performed to infer the taxonomy and functional changes in all the treatment groups. RCJ administration was explored on colon microbial alpha diversity using Shannon indices, a measure of evenness in the samples, on taxonomy analysis results at the species level. The DSS group showed a decrease in alpha diversity (**Fig 5A**). The existing carbohydrates/polysaccharides in RCJ could promote a higher alpha diversity.

**Fig 5:**
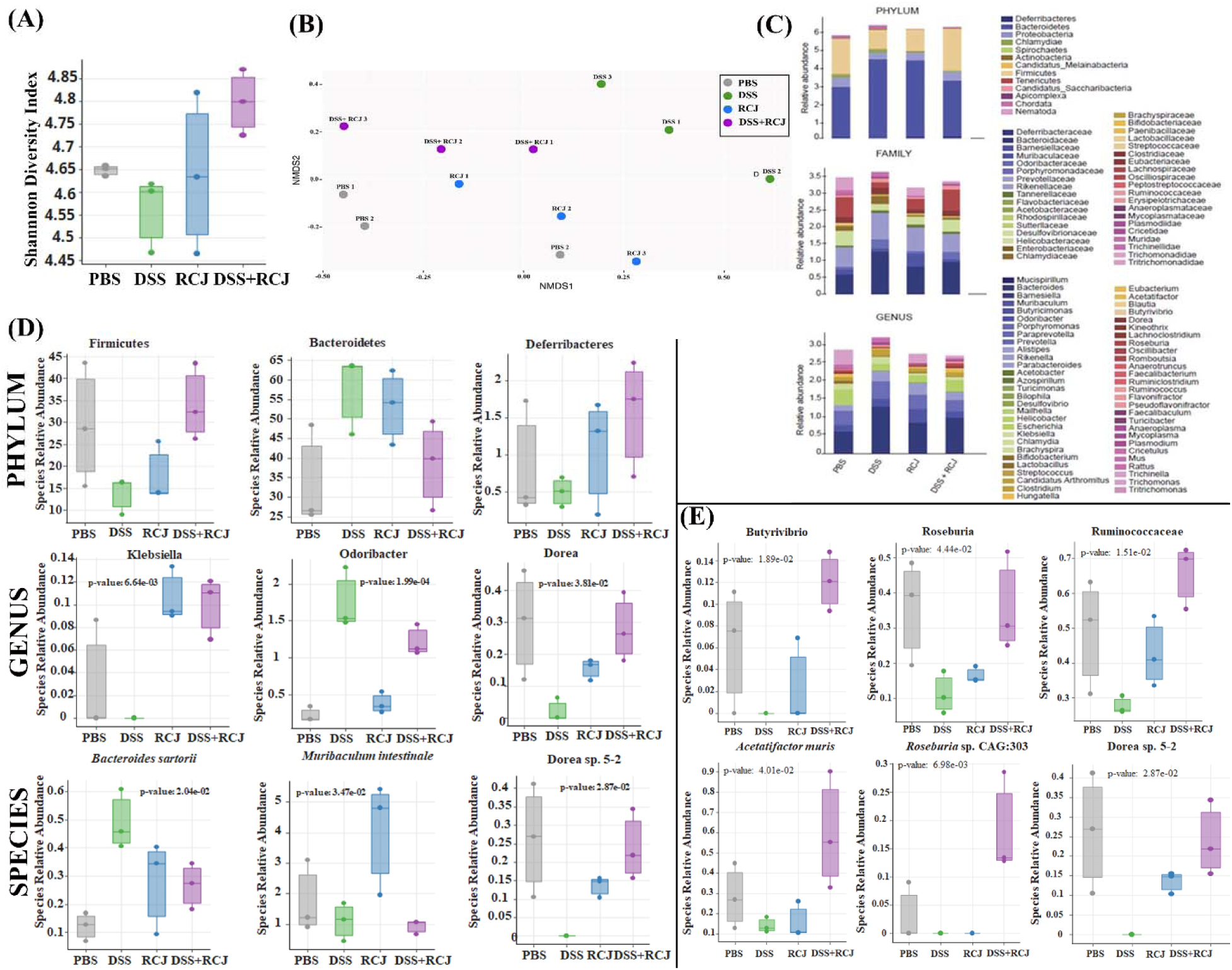
Effect of RCJ treatment on gut microbiota. **(A)** Alpha diversity using Shannon indices, a measure of evenness in sample. (B) NMDS plots representing the samples’ closeness when compared to the control. **(C)** Relative abundance of taxa. **(D)** Relative abundance of significant organisms at phylum, genus, and species level. **(E)** Relative abundance of some reported SCFA-producing taxa. Statistical significance was determined using one-way ANOVA, followed by the Tukey test.

NMDS plots of taxonomy and pathway data consistently revealed that the microbiome in the RCJ+DSS group was more similar to the control samples than that in the DSS group (**Fig 5B**), supporting the hypothesis that under DSS-induced colitis, RCJ partially restores the microbiome closer to a normal healthy microbiome.

#### RCJ intervention alters the SCFA-producing population in the gut microbiota

Impact of RCJ on both cecal mucosa and luminal microbiota composition by MEGAN analysis. At the phylum level, we did not observe a compelling signal of differentially abundant phyla (**Fig 5C**), except for the following comparisons that were close to the significance threshold **(SF2)** increased abundances of *Bacteroidetes, Chlamydiae, Chordata*, and decreased abundance of Firmicutes in DSS treated group of mice compared with PBS, RCJ, and RCJ+DSS groups; increased abundance of *Firmicutes, Deferribacteres* and reduced abundance of *Bacteroidetes* and *Chordata* in the RCJ+DSS group (**Fig. 5D**).

A comprehensive set of significant taxa was provided in **(SF2).** Here, the focus was on the taxa that were found to be more relevant to colitis and were found to be significant.

At the **genus** level, RCJ treatment significantly enriched *Muribaculum*, *Klebsiella*, and *Desulfovibrio* (**Fig 5D; S4A**) (P < 0.05, 0.005, and 0.05, respectively) compared with the control group. Ruminococcus (**Fig S5A)** and Odoribacter (**Fig 5D**) genera were significantly enriched in the DSS group compared to those in the control group (P <0.05, 0.0413, and 0.0005, respectively). In contrast, *Lachnoclostridium* (**Fig S5A)** and *Dorea* (**Fig 5D)** genera were significantly enriched in the RCJ+DSS group compared to those in the DSS-only group (P-values 0.05 and 0.05, respectively). Furthermore, *Bacteroides sartorii* (**Fig 5D**) and *Ruminococcus flavefacien* (**Fig S5A)** species were enriched in the DSS group (P< 0.05, and 0.05, respectively), while *Clostridium sp.* CAG:557 (**Fig S5A)** *and Dorea sp*. 5-2 (**Fig 5D**) were depleted in the DSS group (P<0.005 and 0.05, respectively). *Muribaculum intestinale* (**Fig 5D**) was strongly enriched by RCJ treatment (P< 0.05). **(SF2)** summarizes significant taxa at different taxonomic levels.

Furthermore, LefSe analysis was performed to detect significantly different taxa at different taxonomic levels (**Fig S4)** and to generate a cladogram representing the taxonomic relatedness of the significant taxa. The cladogram showed substantial differences in 106 taxa among the four treatment groups (PBS, RCJ, DSS, and DSS + RCJ) (**Fig SF3**). Red, green, blue, and purple indicate different groups, with the species classification at the phylum level, class, order, family, and genus shown from inside to outside. For instance, the cladogram clearly indicated that DSS treatment affected members of the *Bacteroidales* family. In contrast, the yellow nodes represent species with no significant differences.

Furthermore, a few key taxa, such as *Butyrivibrio, Roseburia, Ruminococcaceae, Acetatifactor muris, Rosburia Sp.* CAG:303*, Dorea Sp.* 5-2 were found. They were more abundant in the DSS+RCJ group than in the DSS group (**Fig 5E**). These families are *Clostridia* and have been reported to produce butyrate (41, 42). These microbial taxa have been reported to be responsible for SCFAs productions (43–45).

NMDS plots for the CPM-normalized abundance of the biochemical pathways detected from the metagenomic data showed a similar trend to that observed in the taxonomy analysis. RCJ restored the functional profiles of the microbiome closer to those of the control condition (**Fig 6A**). Functional analysis of the microbiome revealed several pathways significantly associated with the DSS group. A comprehensive list of associated pathways is provided **(Fig SF5).** Here, several significant pathways were highlighted that were found to be relevant to colitis: arginine synthesis, biotin biosynthesis, long-chain fatty acid biosynthesis (oleate biosynthesis and 5Z-dodec-5-enoate biosynthesis), heme biosynthesis, L-L-histidine degradation, while they were brought back closer to control by the RCJ treatment (**Fig 6B**). On the other hand, arginine synthesis and l-histidine degradation, which were upregulated in the DSS group compared with the control, remained upregulated in the RCJ+DSS group compared with the control group (**Fig 6C**). The pathway for L-glutamate degradation to propionate was one of the very few pathways that showed an enrichment trend only in the RCJ+DSS group (**Fig. 6C**). *Bacteroides sartorii, and Bacteroides caecimuris* are the two main species associated with arginine synthesis and L-histidine degradation. (**Fig 6D**).

**Fig 6:**
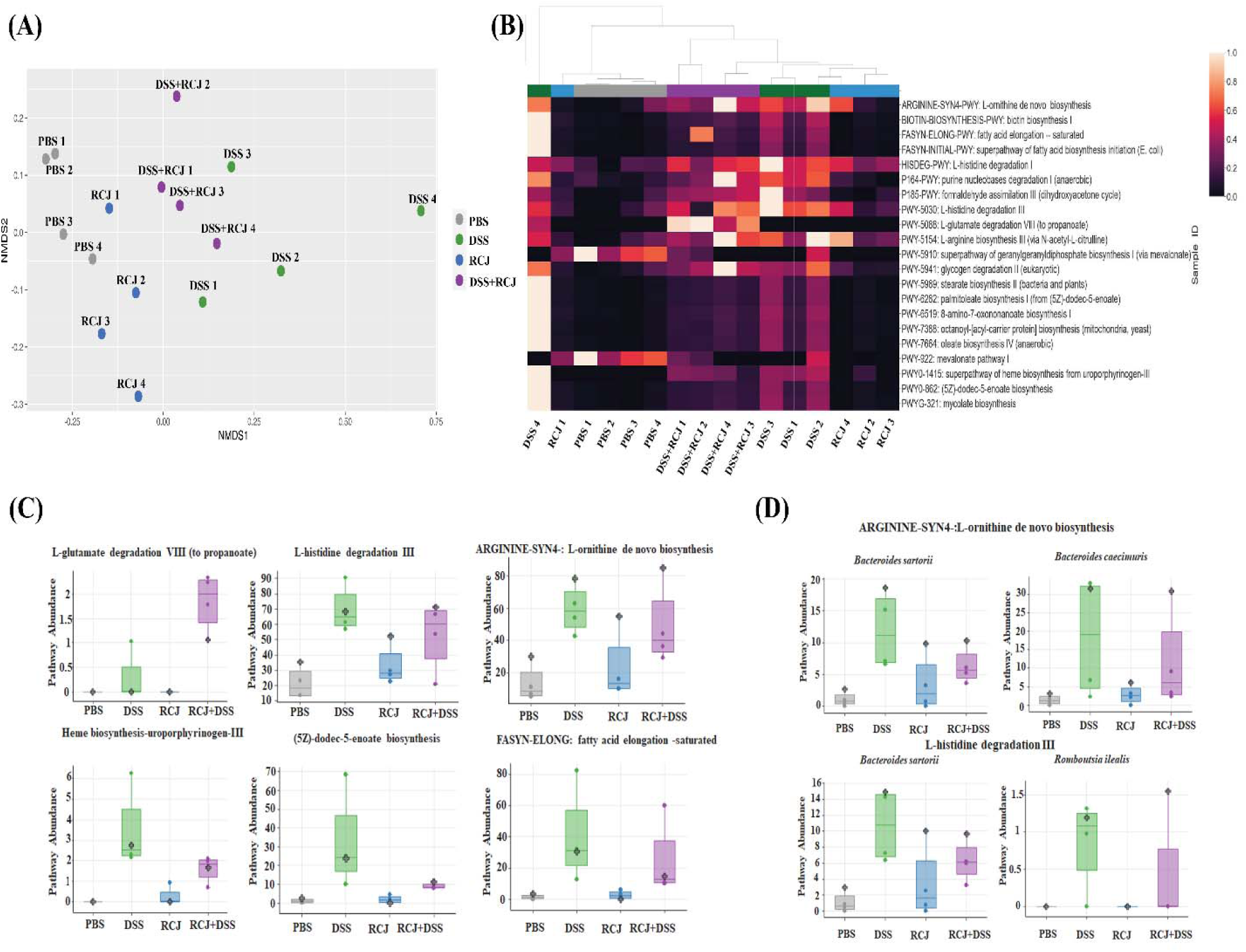
Effect of RCJ on biochemical pathways **(A)** NMDS plots for the biochemical pathways in the microbiota **(B)** Differentially abundant biochemical pathways across treatment groups. **(C)** Top significantly regulated pathways that are beneficial for the colon epithelium health **(D)** Selected organism-specific pathways with significantly differential abundance. *Bacteroides sartorii, and Bacteroides caecimuris* are the two main species associated with arginine synthesis and L-histidine degradation.

#### RCJ, is reversing the dysregulation of immunological responses in DSS-induced colitis mice

The release of pro-inflammatory cytokines was reduced following treatment with RCJ; thus, overall inflammation was induced by DSS. Therefore, we aimed to elucidate the role of immune cells in the regulation of these events. First, we examined two central immune cell populations, macrophages and T cells, stained for F4/80 and CD3 markers. Both populations were increased in the DSS group. Interestingly, the DSS+RCJ both had significantly (P<0.05; P<0.01) fewer macrophages and T cells (**Fig 7A,7B**).

**Fig 7:**
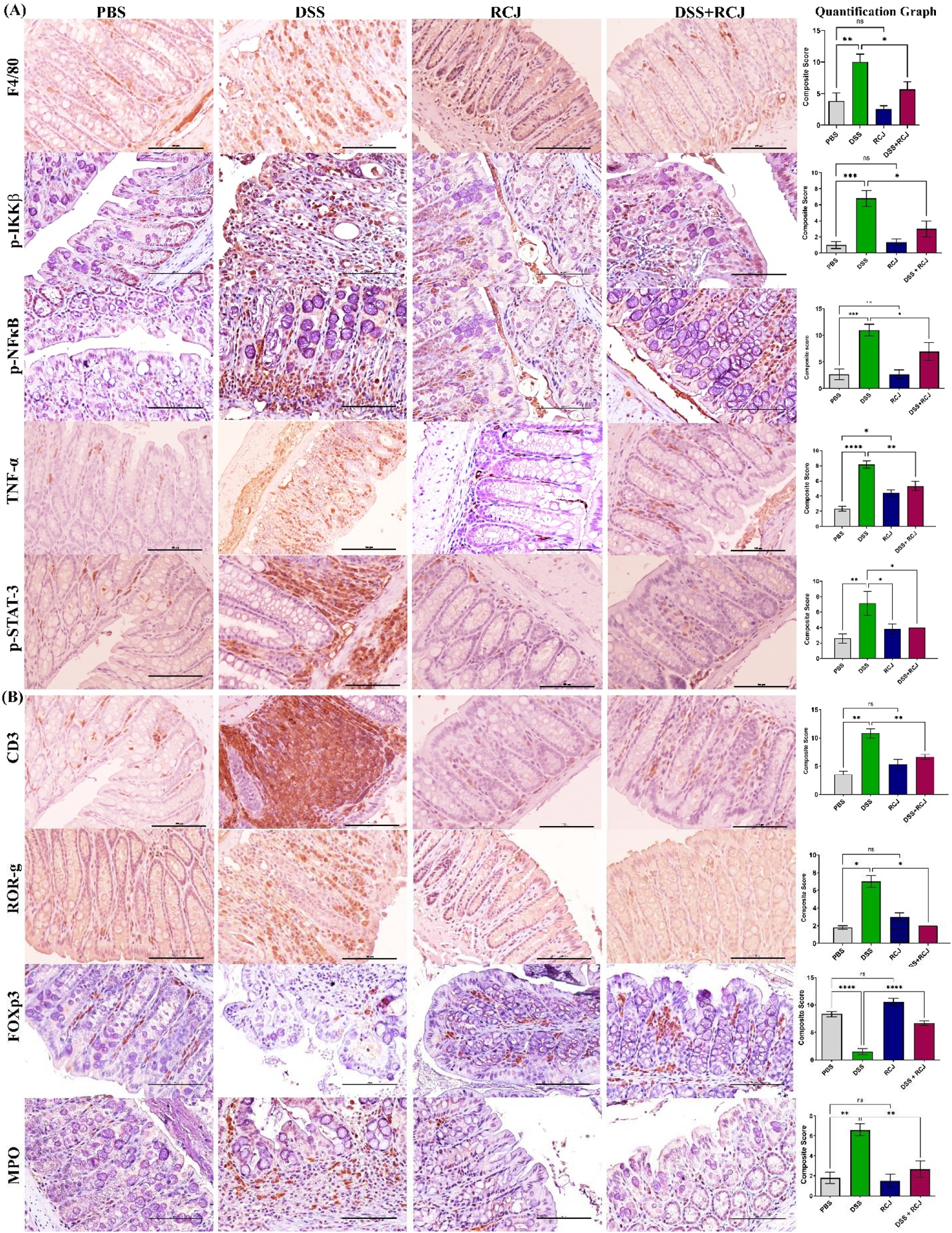
RCJ, reversing the dysregulation of immunological responses in DSS-induced colitis mice **(A)** Depicts IHC staining for F4/80 (total macrophages), and to check inflammation stained for p-IKKβ, p-NF-κB, and p-STAT **(B)** Shows IHC staining for T cell marker panel with CD3, RORγ, FOXp3, and MPO for the neutrophils, All values represent the means ± SD; error bars in the histograms are the standard error of the mean. Scale bars represent 100 µm for the IHC. Statistical significance was determined using one-way ANOVA, followed by the Tukey test. *P ≤ 0.05, **P ≤ 0.01, ***P ≤ 0.001

Due to the loss of intestinal barrier function in DSS-treated animals, the loss of tight junction proteins and epithelial cell damage would lead to leakiness. This would allow microbial translocation (lipopolysaccharides (LPS) from gram-negative bacteria) with the release of other bacterial endotoxins from the colonic lumen to the lamina propria. These events trigger the maturation of T helper 17 (Th17) and recruitment of neutrophils in the lamina propria, which aggravates oxidative stress and secretion of G-CSF, as confirmed by our cytokine array analysis. Additionally, G-CSF stimulates the bone marrow to produce more neutrophils. Thus, ROR-γ, a specific marker for the Th17 T cell subtype, and MPO-specific marker for neutrophils were stained. The DSS group had a higher ROR-γ and MPO-expressing cell population than the DSS+RCJ group (**Fig 7B**), which signified (P<0.05; P<0.01) that RCJ treatment reduced Th17 T cell maturation and neutrophil recruitment.

Previous studies have shown that colonic Foxp3^+^ regulatory T cells (Tregs), an anti-inflammatory subset of CD4^+^ T cells, maintain immune homeostasis (46). Thus, we checked for the Treg population in the colonic region and found that the DSS group had a significantly lower Foxp3^+^ Treg cell population than the DSS+RCJ group, suggesting that RCJ could increase the levels of crucial immune cells and Tregs (P<0.001) (**Fig 7B**) that secrete IL10, an important anti-inflammatory cytokine.

Furthermore, lipopolysaccharide (LPS) can cause inflammation by activating TLRs on macrophages present in the lamina propria. This, in turn, activates the NF-κB pathway, which is a ubiquitous transcription factor that is well characterized and is a primary mediator of the inflammatory response during inflammation and increases pro-inflammatory cytokine levels TNF-α, IL6, IL-1β, and COX2 levels in these macrophages. Increases in TNF-α act as autocrine signaling for the activation of NF-κB, and an increase in IL6 causes the activation of the STAT3 pathway. p-IKKβ, pNF-κB, TNF-α, and pSTAT3 levels were increased in the DSS group, whereas they were significantly reduced in the DSS+RCJ group (P<0.05, P<0.05, P<0.01;<0.01) (**Fig 7A**). Overall, RCJ treatment inhibited the recruitment of immune cells and lowered the degree of inflammation in DSS-induced colitis.

Our data analysis also showed that an increased utes-to-Bacteroidetes (F/B) ratio indicates high SCFAs production in the RCJ+DSS group compared to that in the DSS group (47, 48). This was also evidenced by increased PPAR-γ expression. Microbiota-derived SCFAs in the gut help regulate the host immune response, maintain the intestinal mucosal barrier, and balance intestinal microbiota (49, 50). PPAR-γ is a crucial anti-inflammatory mediator that can sense butyrate and is expressed at high levels in the colon. Butyrate is an important SCFA known to modulate immune response (51). To further determine whether RCJ ameliorates colitis via microbiota-derived SCFAs/PPAR-γ, PPAR-γ expression was investigated in the colon. PPAR-γ levels were dramatically reduced in the DSS group compared to those in the DSS+RCJ group (**Fig 8A & B**).

**Fig 8:**
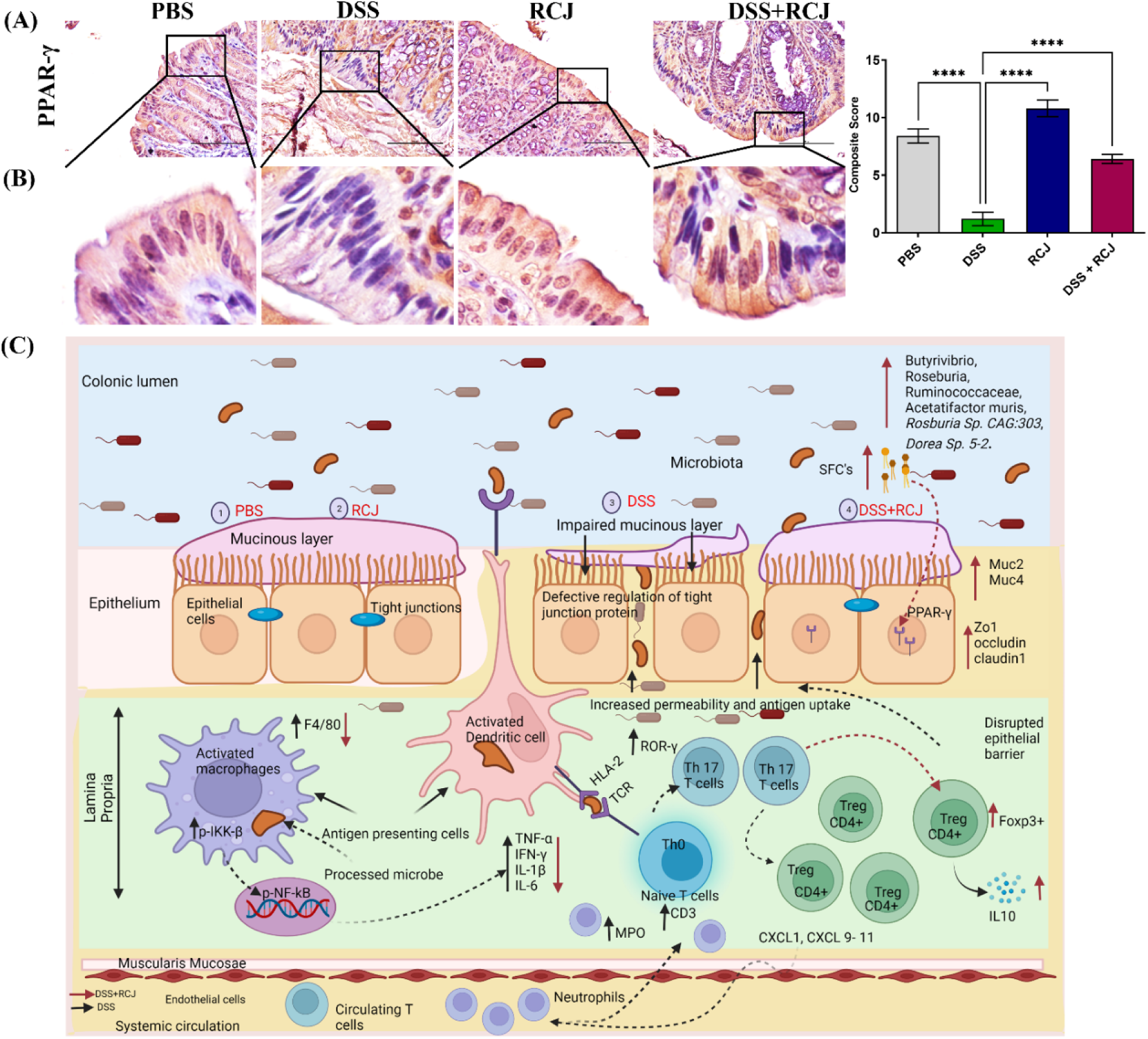
**(A)** Shows IHC staining for anti-inflammatory mediators PPAR-γ indicative of butyrates presence. Scale bars represent 100 µm for the IHC. Statistical significance was determined using one-way ANOVA, followed by the Tukey test. *P ≤ 0.05, **P ≤ 0.01, ***P ≤ 0.001 (**B**) Zoom image of PPAR-γ expression **(C**) A schematic model showing the mechanism by which RCJ alleviated DSS-induced colitis. Intestinal microbiota, oxidative stress, inflammation, and barrier integrity are all affected. RCJ treatment changed the gut microbiota by enriching bacteria such as *Butyrivibrio, Roseburia, Ruminococcaceae, Acetatifactor muris*, *Rosburia Sp. CAG:303*, *Dorea Sp. 5-2*. which subsequently led to increased production of SCFAs such as butyrate, which was evidenced by increased expression of papr-γ leading to a cascade of events anti-oxidative, anti-inflammatory, and barrier-protective responses. Ultimately, intestinal epithelial homeostasis is attenuated, and colitis is attenuated.

## Discussion

Recently, several dietary supplements have emerged as promising therapeutic interventions for IBS and its associated diseases. In the current study, we investigated the effect of RCJ in a DSS-induced colitis mouse model. This model is currently widely used to study IBD because of its resemblance to human ulcerative colitis conditions. IBD in humans is an inflammatory disorder of the gastrointestinal tract that is linked to the composition and function of the intestinal microbiota (52). Consumption of RCJ is inversely correlated with inflammation and oxidative stress owing to a group of bioactive compounds present in RCJ (19),(16),(15). However, the active RCJ components that regulate gut microbiota to confer anti-inflammatory function and intestinal homeostasis are unclear.

Here, we focused on ameliorating colitis by RCJ treatment via gut microbiota modulation. Oral administration of RCJ markedly ameliorated DSS colitis, as demonstrated by body weight loss and higher survival rate. In addition, mice recovered when receiving RCJ during the resting period after the first cycle of DSS. The significant features of the DSS-induced colitis model are short colon length, high blood levels in feces, increased diarrhea, and DAI. Notably, the treatment group that received RCJ followed by DSS treatment did not show a colon length shortening effect, considering that the reduction in colon length is a classic indicator of experimental colitis (53). Our results showed that supplementation with RCJ significantly reversed these critical features of colitis. This is consistent with previous reports where Rhein, *Ilex kudingcha*, and Pu-erh Tea Extract treatment alleviated DSS-induced colitis (52, 54, 55). DAI is an indicator of disease severity and is comparable to the clinical representation of IBD. Treatment with RCJ reduced disease severity in the DSS group, indicating that RCJ might function as a prebiotic to provide a novel and effective colitis prevention and therapy strategy.

To maintain intestinal homeostasis, intestinal barrier function and permeability are vital, as shown by experimental and clinical data (56). Intestinal epithelial cells, the mucus layer, and tight junction proteins mainly contribute to intestinal permeability. H&E staining revealed inflammatory cell infiltration, intestinal architectural changes, loss of villi and necrosis of the intestinal surface, reduced cell proliferation, and increased apoptosis in DSS-treated mice.

Moreover, RCJ significantly attenuated DSS-induced oxidative stress by enhancing the expression of antioxidant enzymes, such as SOD and GPX4 and reducing 4 hydroxynonenal expression, which is a crucial mediator of oxidative stress-induced cell death in the colon, which is corroborated by previous studies (57).

To determine the effects on the protective mucin layer, PAS staining for acid and neutral mucin showed the most significant loss of mucins in the DSS-treated group. H&E staining also showed that the PBS control group displayed intact colonic mucosa, crypts, stroma, and submucosa, with no inflammatory cell infiltration in the submucosa or ulceration. In contrast, the DSS group showed severe epithelial cell damage, shortening and hyperplasia of crypts, submucosal edema, inflammatory lesions, severe inflammatory cellular infiltration in the submucosa, and macroscopic spaces between the crypts. However, RCJ treatment ameliorated the colon inflammatory symptoms and intact surface epithelium, leading to less inflammatory cell infiltration in crypts and only mild submucosal edema, a condition of the colon close to that of PBS mice. PAS and Alcian blue staining showed that the DSS-treated group had a significant reduction in the thickness of colonic epithelial mucosa, which was attenuated to near-normal thickness.

In contrast, other groups retained mucin with increased expression of MUC2 and MUC4, suggesting that RCJ plays a protective role against inflammatory colitis, as determined in a mouse model (58). Mucins are important in protecting the gastrointestinal tract and eliminating bacterial toxins. Recent studies have suggested that mucins initiate inflammatory bowel disease, leading to cancer progression (59).

From our initial H&E data, we found disruption of the crypt structure in the DSS-treated group, which was improved in the DSS+RCJ group. Thus, to understand how RCJ enhances intestinal barrier function, the tight junction proteins Claudin-, Occludin, and ZO-1 were assessed as they play a vital role in maintaining intestinal barrier function. Furthermore, we assessed tight junctions as proteins responsible for regulating cellular permeability. Decreased expression of tight junction proteins is observed in most IBD cases. Specifically, the expressions of Claudin, Occludin, and ZO-1 were examined. Several factors, including dietary components, gut microbiota, and cytokines, regulate intestinal tight junction proteins (60). Previous reports have shown that ZO-1 downregulation may be one of the causes of ineffective mucosal healing in IBD patients (61). The gut microbiota modulates the immune system by releasing several mediators such as short-chain fatty acids (SCFAs). These mediators, which are released by immune cells, can induce tight junctions dysfunction. Our results also showed that supplementation with RCJ enhanced the expression of tight junction proteins. This finding aligns with a previous report where ZO-1 expression was improved in mouse intestinal epithelial cells when treated with hyaluronan (62).

Immune cells secrete cytokines that are small peptides. Cytokines act as vital pathophysiological regulators that govern the occurrence and development, ofIBDand, ultimately lead to inflammation as a characteristic feature of IBD (22). The innate immune response plays a pivotal role in IBD, and DSS-mediated stimulation of pro-inflammatory cytokines is involved in colitis (63). The current study demonstrated that DSS supported this trend by increasing pro-inflammatory cytokines, while RCJ administration attenuated this inflammatory response. RCJ alleviated colonic inflammation by reducing inflammatory cytokines, such as TNF-α, IL-6, IL-1β, and IFN-γ, and increasing IL-10 in the plasma of DSS-treated mice. Proinflammatory cytokine genes have a binding site for NF-κB and regulate transcription (64). In addition, DSS treatment causes damage to the colon epithelial cells and leads to tissue inflammation due to the accumulation of nitric oxide (NO) generated by iNOS. RCJ treatment reduced iNOS expression and alleviated inflammatory effects.

Furthermore, the elevated COX-2 in DSS was due to loss of gut permeability, as the cell wall component LPS of gram-negative bacteria can stimulate epithelial cells to promote the production of COX-2 and cause inflammation (65), which was reversed by RCJ supplementation. The RCJ-treated group showed reduced macrophage (F4/80) and increased T-reg (Foxp3) cell infiltration, crypt destruction, and ulcer formation due to reduced pro-inflammatory cytokines (such as TNF-α, IL-6, IL-1β, and IFN-γ) and increased anti-inflammatory IL-10 in serum. These results are highly consistent with previous reports (66).

Gut microbiota contributes to immune responses and intestinal permeability through a variety of enzymes and metabolites. This study found that DSS treatment results in microbiome shifts from both taxonomic and functional perspectives, and that RCJ can be used to mitigate this change. The low abundance of *Firmicutes* and high abundance of *Bacteroidetes*, a signature of microbiota dysbiosis in DSS, where RCJ supplementation reversed these effects.

Nutraceuticals are attracting increasing attention worldwide as potent therapeutic agents for IBD owing to their fewer side effects as they contain different bioactive compounds. It is an opportunity for direct contact between the compounds and colon epithelial cells and indicates a close relationship with gut microbiota alteration when supplemented (67). Such interactions between active compounds and the gut microbiota include 1. direct modulation of the gut microbiota by bioactive compounds in the RCJ, 2. specific bacterial strains that transformed the bioactive compounds of RCJ, and 3. Bioactive compounds in RCJ regulate the metabolism of the gut microbiota (68)

SCFAs (acetate, butyrate, and propionate) are produced in the colonic lumen, mainly by the fermentation of dietary fiber by the gut microbiota. These SCFAs are vital for physiological and pathophysiological effects on colonic events (69). For example, SCFAs are known to alleviate colitis conditions by reducing the production of pro-inflammatory cytokines, thus blocking the NF-κB and STAT 3 signaling pathways and, hence, reducing colon inflammation.

Taxonomic analysis of our metagenomic data suggests that several taxa known for SCFAs production activity were enhanced in the DSS+RCJ treatment groups. These SCFAs act as a significant energy source for colonic mucosal cells and are essential regulators of gene expression during crucial events such as apoptosis, differentiation, and inflammation. These microbiota-derived SCFAs prevent infections by modulating the systemic immune response and enhancing the intestinal mucosal immune barrier (70). Thus, SCFA-producing bacteria act as probiotics and play vital roles in various biological functions.

Butyric acid-producing genera that reside in the human intestinal tract and their underrepresentation have been linked to disease states (43, 71–73). Our data showed that these populations were destroyed by DSS treatment, while RCJ effectively recovered them in the RCJ and RCJ+DSS groups. More specifically, *Dorea sp.* 5-2 species stands out as it has an undetectable population in the DSS group, while its abundance in RCJ and RCJ+DSS was closer to that in the control group. This suggests an interesting hypothesis about the mechanism by which RCJ improves the microbiome and inflammation status by enriching this specific microbial pathway related to SCFA production in an unhealthy microbiome. In contrast, DSS treatment promoted the population of *Bacteroides sartorii*. Our pathway analysis data indicated *Bacteroides sartorii’s* role in increasing the abundance of the arginine biosynthesis pathway. This suggests that *Bacteroides sartorii* caused an imbalance in arginine levels in the DSS group.

As high levels of butyrate-producing bacteria were found in the RCJ+DSS group, PPAR-γ expression was determined to be an anti-inflammatory mediator, and butyrate levels in the colon controlled its expression. Under DSS conditions, PPAR-γ levels were significantly low. To further validate the effect of RCJ on the colitis model via microbiota-derived SCFAs/PPAR-γ, it was found that the RCJ+DSS treatment group showed increased expression, suggesting that IBD is correlated with PPAR-γ deficiency. This trend was described in previous reports where phloretin enhanced PPAR-γ expression in DSS-induced mice (74). PPARγ is also known to inhibit the nuclear factor-κB (NF-κB) and MAPK pathways. Regulation of these key signaling networks inhibits the production of cytokines and chemokines, which reduces the influx of inflammatory cells into inflamed tissues (75). Based on our current findings, it can be concluded that SCFAs derived from microbiota ameliorated colitis in mice by increasing PPAR-γ expression. Butyrate may also affect the histone acetylation of gut CD4+ T cells to epigenetically control the production of genes necessary for Treg cell induction. Previous studies have shown that colonic Foxp3^+^ regulatory T cells (Tregs), which are an anti-inflammatory subset of CD4^+^ T cells, maintain immune homeostasis (46). Our results showed that the Foxp3^+^ Treg cell population was increased in the DSS+RCJ group, which might be crucial for immune cells that secrete IL10, a vital anti-inflammatory cytokine (76).

A mechanistic understanding of dietary input and gut microbial metabolism is important because microbiome-derived metabolites lead to divergent physiology and help in gut homeostasis (77). The overall trend of the microbial metabolic pathways that were significantly detected showed that RCJ restored the imbalance introduced by DSS. However, it is not possible to directly infer whether the microbiome is only affected by DSS treatment or whether DSS plays a role in initiating or influencing pathology. An increasing number of studies indicate the importance of fatty acids as pro- or anti-inflammatory agents (78, 79). Early metabolomics studies on IBD have determined increased levels of long-chain fatty acids in patients with IBD (80). Specifically, octanoate and decanoate have been shown to accumulate in considerable amounts in tissues under pathological conditions, leading to impairment of the functioning of mitochondrial respiratory complexes (81), which is the opposite of the supposed role of short-chain fatty acids as a readily available source for epithelial cells (82). The elevated level of reads mapped to the microbial biosynthesis of LCFAs in the DSS-treated group. Interestingly, glutamate degradation to propionate was elevated in the DSS + RCJ group. This indicates an increase in the SCFA producer population. This evidence suggests a link between colitis and microbiome metabolism. During colitis, the production of LCFAs is favored, while treatment with RCJ favors SCFA over LCFAs. The reason for this selective behavior is unclear. Oxidative stress induced by DSS treatment could be an initiator that shifts microbiome metabolism. This is intensified when DSS treatment and induced colitis mucosal permeability are compromised, providing a higher concentration of available oxygen (83). The higher abundance of heme biosynthesis pathways in the DSS group also supports this hypothesis. It is a well-known antioxidant. This suggests that reducing oxidative stress by RCJ treatment might be an important mechanism for the regulation of gut microbiome metabolism. The observed increase in glutamate degradation for propionate production in RCJ-supplemented mice also suggests a metabolic change with RCJ treatment after DSS treatment. DSS treatment also caused the microbial metabolism of several amino acids to shift compared with that in the control group. Arginine biosynthesis genes were present in higher abundance in the DSS group. Higher concentrations of arginine can produce nitrite oxide, which is increased in patients with UC (84). The same trend was observed for L-histidine degradation pathways. More histidine degradation by the bacteria means that a larger sink for L-histidine in the intestinal tract can result in lower than normal levels in the DSS treatment group. A lower available histidine concentration is associated with a higher risk of relapse in UC patients (85).

In conclusion, DSS treatment depleted the population of SCFA-producing bacteria and promoted the population of pathogenic bacteri*a.* Furthermore, this population shift is accompanied by dysregulation of the gut microbiome metabolism, especially fatty acid and amino acid metabolism. Verifying the observed taxonomic trend from a functional perspective requires further metagenomic studies coupled with meta-transcriptomic datasets. Further, there is a need to identify critical active ingredients of RCJ that enhance these beneficial bacteria, which could thus be used as a promising clinical therapy. However, the exact molecular mechanisms underlying SCFAs need to be delineated.

## Conclusion

In summary, RCJ attenuated DSS-induced colitis in mice by altering the gut microbiota and enriching SCFA-producing bacteria, such as *Butyrivibrio, Ruminococcaceae, Acetatifactor muris*, *Rosburia Sp.* CAG:303, *Dorea Sp.* 5-2. The pathway abundance showed proof of SCFAs (L-glutamate degradation to propionates). Bacteria such as *Bacteroides sartorii, and Bacteroides caecimuris,* responsible for histidine degradation, significantly reduced the RCJ+DSS-treated group. Further increased expression of PPAR-γ was found in RCJ-treated groups, which evidences the presence of SCFAs (butyrate). PPAR-γ inhibits the activation of the NF-κB signaling pathway and reduces the expression of IL6, TNF-α, and iNOS, which are crucial for inflammation. Further, RCJ treatment also increased the Treg cell (FOXP3 ^+^) population as critical immune cells, corroborating the increased IL10 expression, a critical anti-inflammatory cytokine. Thus, these changes in the gut microbiota subsequently led to pronounced gut barrier function, colon repair, and anti-oxidative effects, resulting in the attenuation of intestinal damage and colonic inflammation (**Fig 8C)**. These findings provide novel insights into how RCJ ameliorates DSS colitis by modulating the gut microbiota, which could accelerate the development of preventive and therapeutic strategies for IBD patients.

## Abbreviation

RCJ: Red Cabbage juice
IBD: inflammatory bowel disease
UC: ulcerative colitis
DSS: dextran sulfate sodium
CDC: Centers for Disease Control and Prevention
SCFAs: Short-chain fatty acids
LCFAs: long-chain fatty acids
TNF-α: Tumor necrosis factor alpha
GSLs: Glucosinolates
PEF: pulsed electric field
ITCs: isothiocyanates
UV: ultraviolet
HPLC: High-performance liquid chromatography
DAD: diode-array detector
FLD: fluorescence detector
RID: refractive index detector
IL (1-17): interleukin (1-17)
IFN-γ: interferon gamma
TUNEL: Terminal deoxynucleotidyl transferase dUTP nick end labeling
HRP: streptavidin-horseradish peroxidase
Ppar γ: peroxisome proliferator-activated receptor gamma
iNOS: inducible nitric oxide synthase
COX-2: cyclooxygenase-2
DAPI: 4’,6-diamidino-2-phenylindole
MUC 2: Mucin 2
MUC 4: mucin 4
pSTAT3: phospsho Signal transducers and activators of transcription 3
RORγ: Retineic-acid-receptor-related orphan nuclear receptor gamma
MPO: Myeloperoxidase
LDA: linear discriminant analysis
LEfSe: linear discriminant analysis effect size
STAMP: Statistical Analysis of Metagenomic Profiles
CPM: Count per million
MaAsLin: Multivariable association discovery in population-scale meta-omics studies
NMDS: Non-metric multidimensional scaling
GC-MS: Gas chromatography mass spectrometry
SOD: superoxide dismutase
4-OH-enol: Hydroxynonenal
GPX4: glutathione peroxidase 4
CXCL: C-X-C motif chemokine
Th17: T helper 17
UC: ulcerative colitis
NF-κB: nuclear factor kappa-B
MAPK: mitogen-activated protein kinase
IKKβ: inhibitor of nuclear factor kappa-B kinase
G-CSF: granulocyte colony-stimulating factor
GM-CSF: granulocyte-macrophage colony-stimulating factor
CD3: Cluster of differentiation 3
LPS: Lipopolysaccharide
TLRs: Toll-Like Receptors
DAI: disease activity index
Tregs: T regulatory cells
Th17: T helper 17
IKKβ: inhibitor of nuclear factor kappa-B kinase

## Supplementary Material

Additional File 1: Supplementary table 1.

Additional File 2: Supplementary table 2.

Additional File 3: Supplementary figure 1.

Additional File 4: Supplementary table 3.

Additional File 5: Supplementary figure 2.

Additional File 6: Supplementary figure 3.

Additional File 7: Supplementary figure 4.

Additional File 8: Supplementary figure 5.

## Acknowledgments

The authors thank the Complex Carbohydrate Research Center at the University of Georgia, Georgia, USA., for their support. The authors thank the Biorender web tool for generating animated figures, the DNA core UNMC, and the bioinformatic core UNMC. This work was supported by the National Science Foundation (awards ACI-1532235 and ACI-1532236), University of Colorado Boulder, and Colorado State University. The Summit supercomputer is a joint effort of the University of Colorado Boulder and the Colorado State University.

## Author’s Contribution

E. J. B. performed mouse studies and reviews; N.S.N. Conceptualization, experimentation, writing the original draft, review, and editing; P.G., S.P., S.H.J.C., - Shortgun metagenomic sequencing analysis R.P.R., M.P.S., J.T.K., J.R.D., J.B., C.L.L., J.M.F., S.K.B., J.S., - review and editing; D.R.P – Histopathology analysis; T.M. - oligosaccharides analysis in RCJ; A.A.W. – Total phenolic compound analysis of RCJ. S.R. Conceptualization, review, and editing. All authors have read and agreed to the published version of the manuscript.

## Funding

National Science Foundation (NSF), Nebraska Research Initiative (NRI) Grant, and National Institute of Health (NIH) R01 CA247763.

## Declarations

### Ethics Committee Approval and Patient Consent

All experimental protocols were approved by the Animal Care and Use Committee of the University of Nebraska Medical Center in accordance with the Guidelines for the Care and Use of Laboratory Animals (ICAUC 16-067-10-FC).

### Consent for publication

Not Applicable

### Availability of data and material

The data reported in this paper are accessible in the NCBI Short Read Archive (SRA) under accession ID PRJNA944265 (https://www.ncbi.nlm.nih.gov/sra/?term=PRJNA944265). The original R scripts and data used for statistical analysis are available at GitHub (https://github.com/chan-csu/RCJ_Megtagenomics).

### Competing interests

Christian L Lorson is the co-founder and CSO of Shift Pharmaceuticals, and Surinder Batra is one of the co-founders of Sanguine Diagnostics and Therapeutics, Inc., located in Omaha, NE, USA.

### Authors’ information (ORCID)

Nagabhishek Sirpu Natesh https://orcid.org/0000-0003-3826-1591

Emily Jean Wilson https://orcid.org/0009-0008-9786-1566

Satyanarayana Rachagani https://orcid.org/0000-0003-3949-8566

Surinder K. Batra https://orcid.org/0000-0001-9470-9317

Jussuf T. Kaifi https://orcid.org/0000-0003-3425-2356

Christian L Lorson: https://orcid.org/0000-0002-1023-2169

Marudu Pandiyan https://orcid.org/0000-0002-5217-2950

---

**S Table 1:** Bioactive compounds’ retention during freezing and PEF treatment. **(A)** Total phenolic compounds and free anthocyanins content by spectrophotometry

| Sample ID | pH | Brix | Glucose (g/L) | Fructose (g/L) | Citric acid (g/L) | Malic acid (g/L) | Unknown acid (g/L as malic acid eq.) |
| --- | --- | --- | --- | --- | --- | --- | --- |
| RCJ | 6.42 ± 0.05 | 6.4 ± 0.2 | 18.2 ± 0.2 | 15.1 ± 0.2 | 0.8 ± 0.1 | 3.0 ± 0.3 | 34.2 ± 2.1 |
| Total phenolic compounds and free anthocyanins content by spectrophotometry |  |  |  |  |  |  |  |
| Sample ID | Total phenolic compounds content (mg/L catechin eq.) |  |  | Free anthocyanins content (mg/L malvidin-3-glucoside eq.) |  |  |  |
| RCJ | 382.5 ± 93.5 | | | 257.9 ± 3.1 | | | |

**S Table 2:** HPLC-DAD method to determine the composition of monomeric polyphenols present in the RCJ. *The red cabbage juice was rich in anthocyanins and identifying all the anthocyanins was established only based on the literature review and UV/vis spectrum. Therefore, the name anthocyanins has to be taken with caution.

**S Table 1:** Bioactive compounds' retention during freezing and PEF treatment (A) Total phenolic compounds and free anthocyanins content by spectrophotometry
| Sample | Free anthocyanins content mg/L Cyanidin-3-glucoside eq. | Total hydroxycinnamic acids content (mg/L caffeic acid eq.) |  |
| --- | --- | --- | --- |
| Red cabbage juice | 254.3 ± 23.6 | 55.6 ± 2.3 |  |
| <u>Hypothesized anthocyanins in juice (based on literature and UV/visible spectrum)</u> |  | <u>% of total anthocyanins</u> | <u>mg/L</u> |
| Cyanidin-3-diglucoside-5-glucoside |  | 22.7 | 57.61 |
| Cyanidin-3-(sinapoyl)-diglucoside-5-glucoside |  | 2.5 | 6.34 |
| Cyanidin-3-glucoside-5-glucoside |  | 5.5 | 13.97 |
| Cyanidin-3-(feruloyl)(feruloyl)-triglucoside-5-glucoside |  | 1.9 | 4.88 |
| Cyanidin-3-(sinapoyl)-triglucoside-5-glucoside |  | 0.8 | 2.12 |
| Cyanidin-3-(feruloyl)-diglucoside-5-glucoside |  | 43.6 | 110.78 |
| Cyanidin-3-(feruloyl)(feruloyl)-triglucoside-5-glucoside |  | 3.3 | 8.38 |
| Cyanidin-3-(sinapoyl)(sinapoyl)-diglucoside-5-glucoside |  | 19.7 | 50.23 |

**S Table 3.** Calculated Mol % of monosaccharides and Total Carbohydrate percentage by weight for the sample.

| Glycosyl residue | Cabbage Juice |  |
| --- | --- | --- |
|  | Mass (µg) | Mol %<sup>1</sup> |
| Arabinose (Ara) | 1.6 | 2.4 |
| Ribose (Rib) | 0.6 | 0.9 |
| Rhamnose (Rha) | 0.1 | 0.2 |
| Fucose (Fuc) | 0.08 | 0.1 |
| Xylose (Xyl) | 0.3 | 0.4 |
| Mannose (Man) | 1.0 | 1.3 |
| Galactose (Gal) | 3.5 | 4.5 |
| Glucose (Glc) | 70.4 | 90.1 |
| SUM | 77.6 | 99.9 |
| Total Carbohydrate % by weight | 19.4% |  |

<sup>1</sup>Values are expressed as mole percent of total carbohydrate. The total Mol % may not add to exactly 100% due to rounding.

**S Fig 1.**
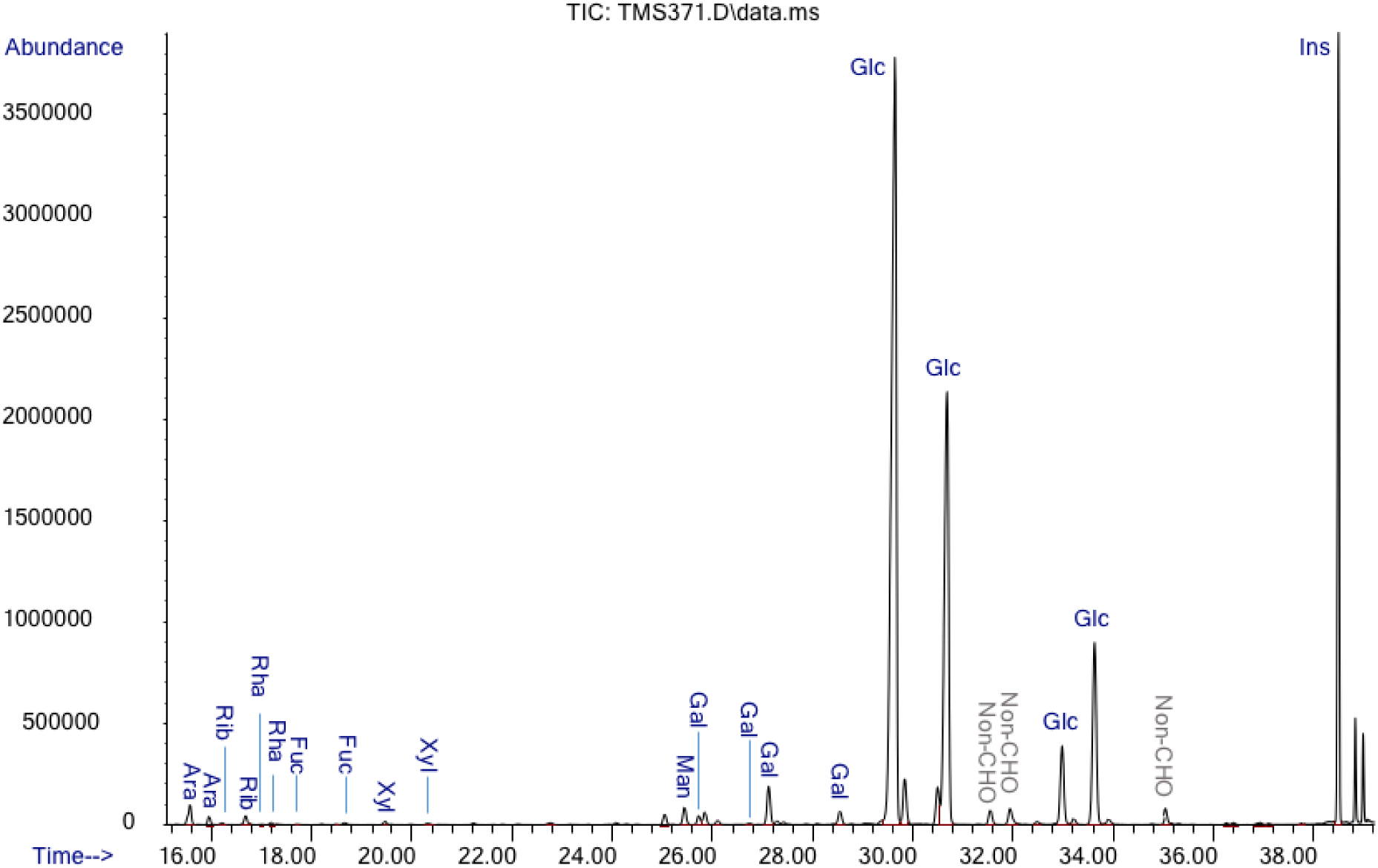
Glycosyl composition analysis by GC-MS of TMS derivatives of methyl glycosides. Shows Total ion chromatogram for the TMS residues from the cabbage juice. “Non-CHO” = non-carbohydrate.

**S Fig 2:**
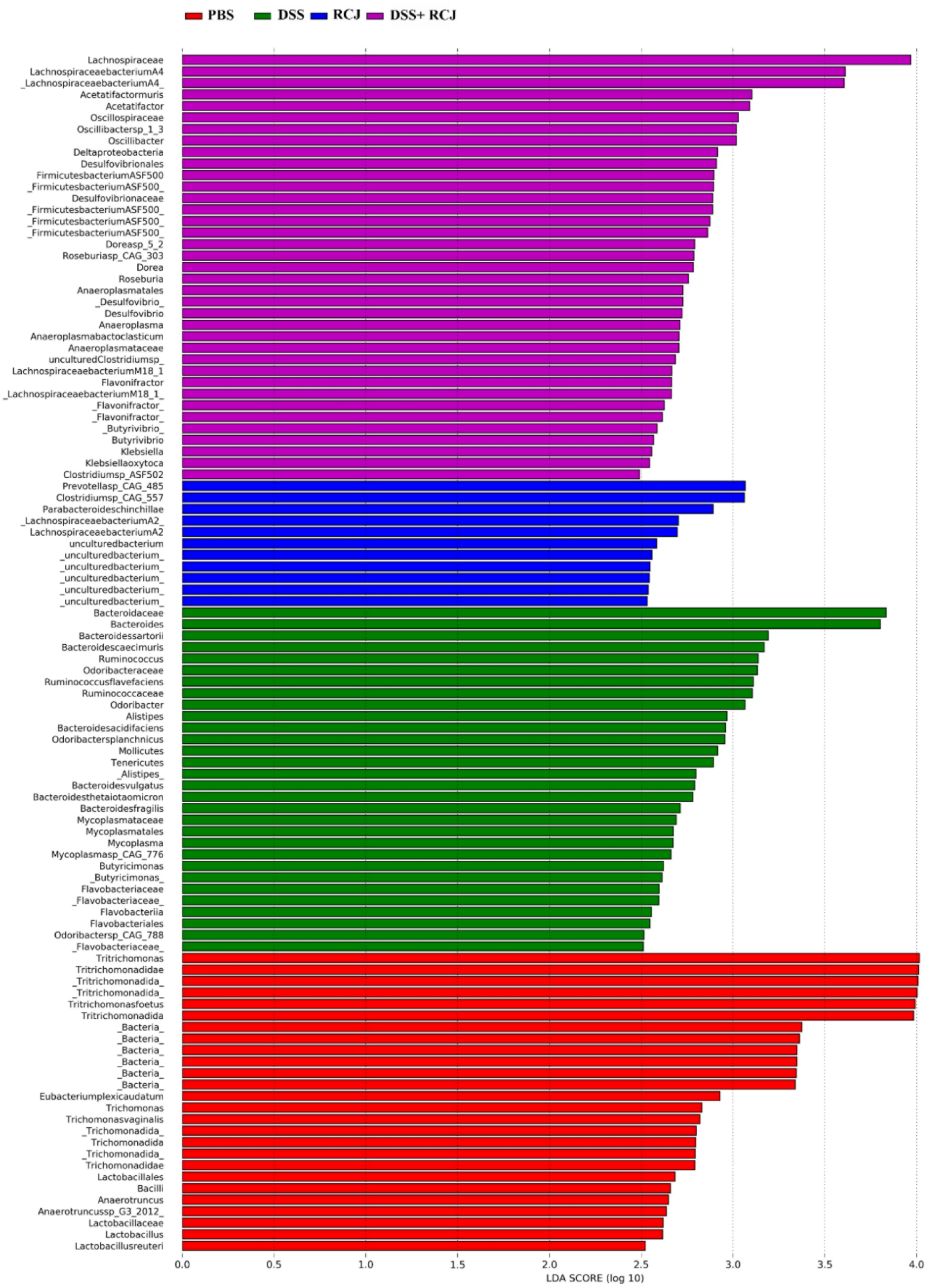
Shows **LefSe** analysis to detect significantly different taxa at the different taxonomic levels

**S Fig 3:**
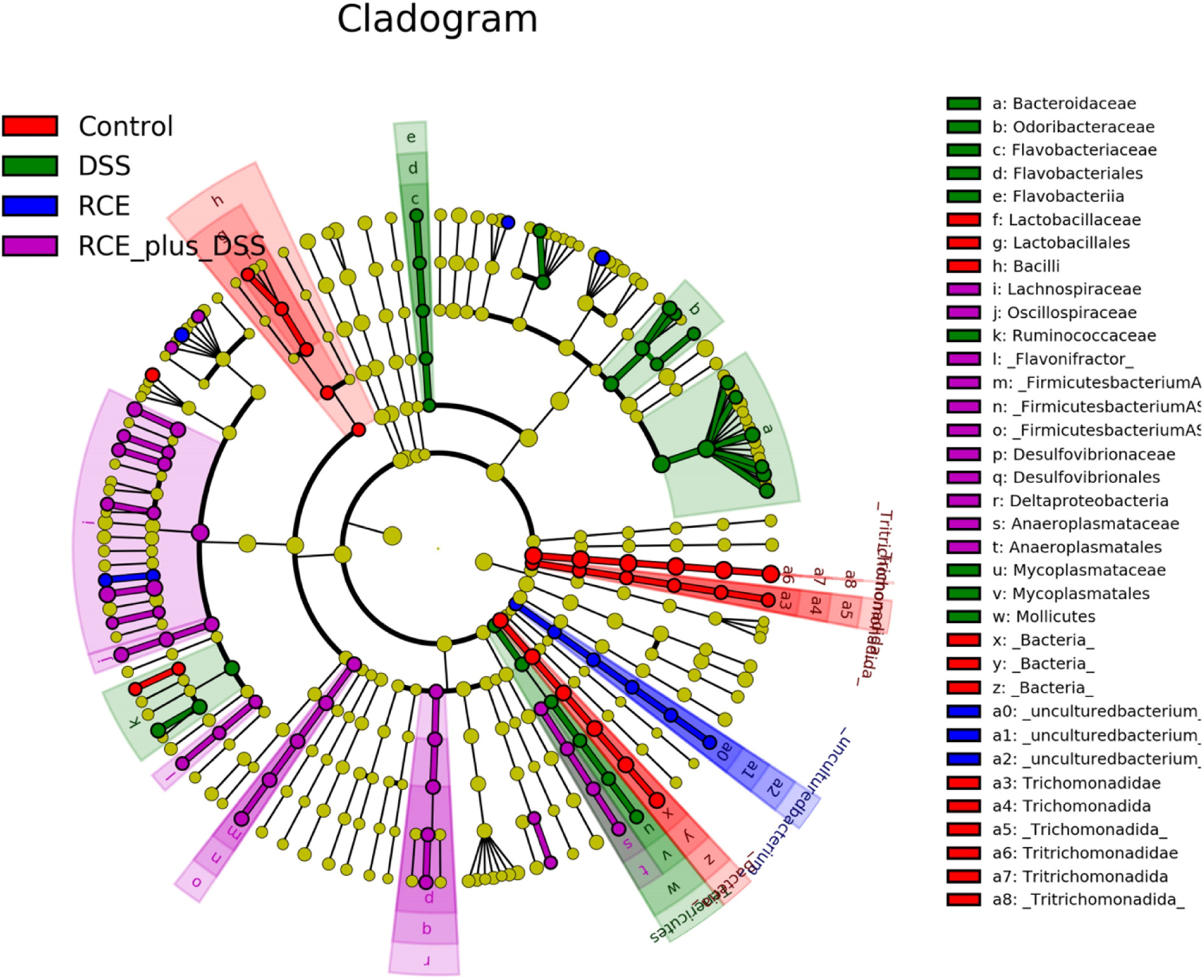
The cladogram showed substantial differences in 106 taxa among four treatment groups (PBS, RCJ, DSS, and DSS+RCJ) Red, green, blue, and purple indicate different groups, with the species classification at the phylum level, class, order, family, and genus shown from the inside to the outside

**S Fig 4:**
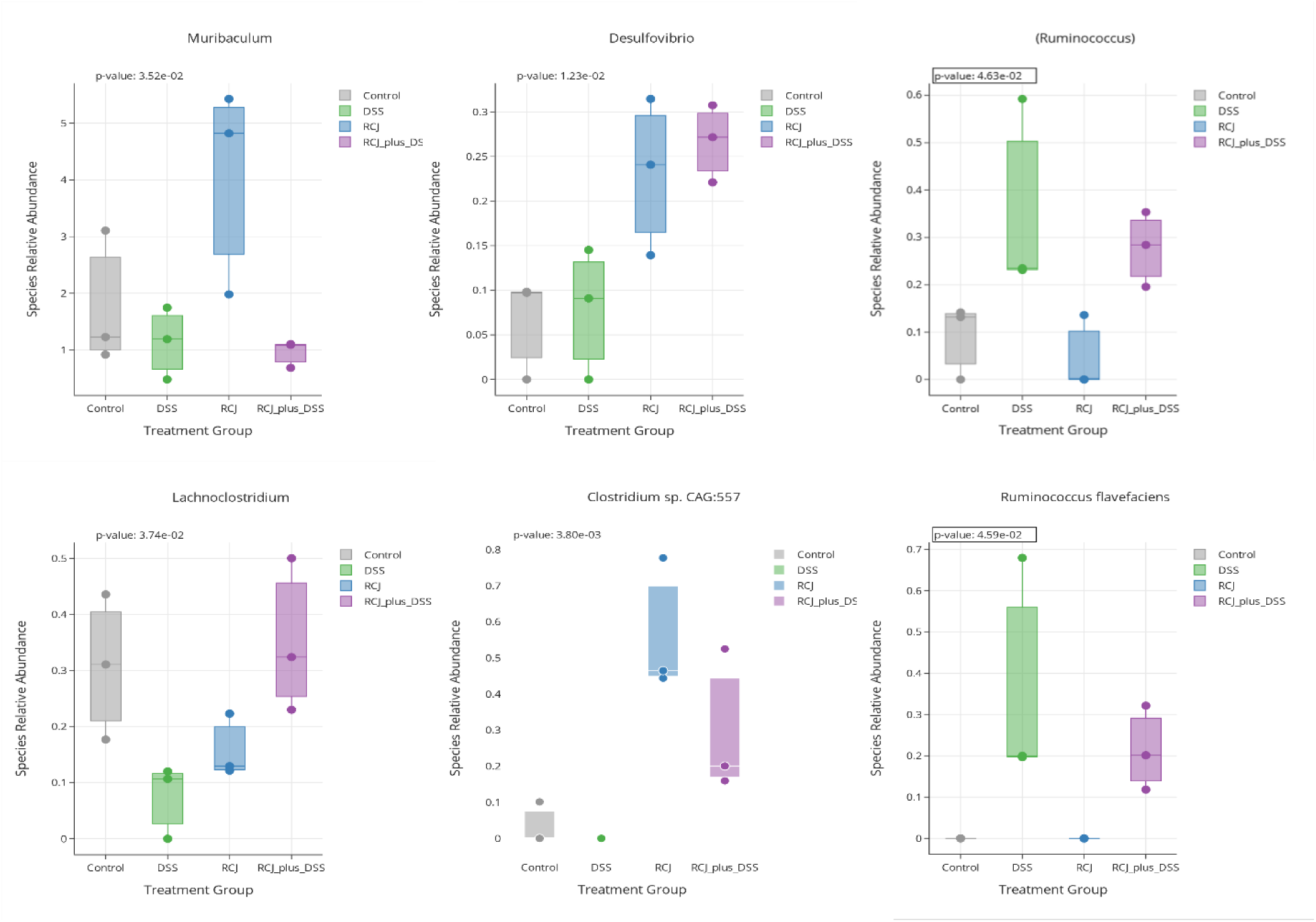
Graphs show the relative abundance of significant organisms at phylum, genus, and species level

**S Fig 5 A & B:**
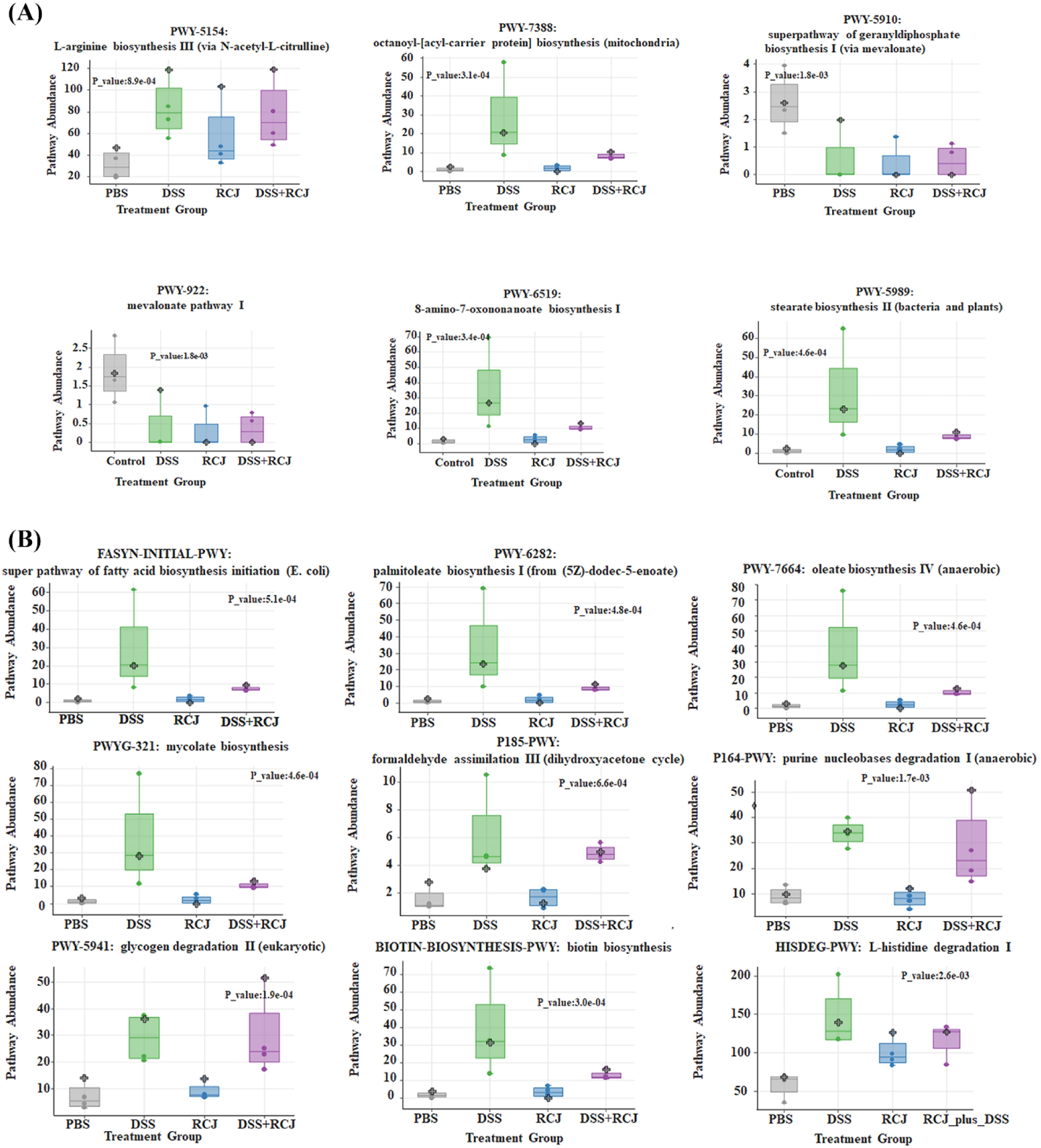
Indicates the rest of the top significantly regulated pathways beneficial for colon epithelium health.

